# ASY3 has dosage-dependent diverse effects on meiotic crossover formation

**DOI:** 10.1101/2024.01.09.574930

**Authors:** Lei Chu, Jixin Zhuang, Miaowei Geng, Yashi Zhang, Chunyu Zhang, Arp Schnittger, Bin Yi, Chao Yang

## Abstract

Crossovers create genetic diversity and are required for equal chromosome segregation during meiosis. Their number and distribution are highly regulated by different, yet not fully understood mechanisms including crossover interference. Crucial for crossover formation is the chromosome axis. Here, we explore the function of the axis protein ASY3. To this end, we use the allotetraploid species *Brassica napus* and due to its polyploid nature, this system allows a fine-grained dissection of the dosage of meiotic regulators. The simultaneous mutation of all four *ASY3* alleles results in defective synapsis and drastic reduction of crossovers, which is largely rescued by the presence of only one functional *ASY3* allele. Crucially, while the number of class I crossovers in mutants with two functional *ASY3* alleles is comparable to that in wildtype, this number is significantly increased in mutants with only one functional *ASY3* allele, indicating that reducing the dosage of ASY3 increases crossover formation. Moreover, the class I crossovers on each bivalent in mutants with one functional *ASY3* allele follow a random distribution, indicating compromised crossover interference. These results reveal the dosage-dependent distinct effects of ASY3 on crossover formation, and provide insights into the role of chromosome axis in patterning recombination.

## Introduction

Crop breeding relies on selecting elite varieties that harbour desirable combinations of genetic alleles. The combination of these alleles is accomplished in meiosis through a new assortment of maternal and paternal chromosomes as well as an exchange of segments of the parental chromosomes through crossovers (COs). However, COs are limited and, especially in crops, not equally distributed, leading to linkage drags that limit the success of breeding schemes. Thus, understanding CO patterning mechanisms is a long-standing goal in plant biotechnology.

COs are also crucial for accurate chromosome segregation during meiosis I. Each pair of homologous chromosomes (homologs) needs to form at least one crossover, the so-called obligate CO, controlled by a mechanism known as CO assurance. In addition, class I COs are subject to a tight regulation that prevents COs from occurring close to each other, a phenomenon called CO interference, which contributes to linkage drags. Limiting breeding efforts, the underlying molecular mechanisms leading to CO assurance and interference are not fully understood.

COs are generated through the repair of programmed DNA double-strand breaks (DSBs) via either of two pathways catalyzed by distinct recombination machineries: one machinery depends on the group of meiosis-specific ZMM proteins (Zip1-4, MSH4-5, and Mer3 in *Saccharomyces cerevisiae*, SHOC1/Zip2, HEI10/Zip3, HEIP1, PTD/Spo16, ZIP4, MSH4-5, and MER3 in *Arabidopsis thaliana*) that catalyzes class I COs, representing the major class of COs (85% to 90% in *Arabidopsis*, 70%-90% in rice, 75%-90% in maize); the other machinery relies on structure-specific nucleases, e.g., MUS81 and FANCD2 (a homolog of Fanconi Anemia Complementation Group D2) in *Arabidopsis*, and promotes the remaining, so-called class II COs, which are not limited by interference (Kurzbauer *et al*, 2017; Thangavel *et al*, 2023; Mercier *et al*, 2015; Zhang *et al*, 2014a; Singh *et al*, 2023; Li *et al*, 2018).

CO formation depends highly on the chromosome axis, a proteinaceous structure assembled along the entire length of each pair of sister chromatids at early meiosis that later transforms into the lateral element of the synaptonemal complex (SC) (Hunter, 2015; Wang & Copenhaver, 2018; Blat *et al*, 2002; Panizza *et al*, 2011; Zickler & Kleckner, 1999, 2015; Ito & Shinohara, 2023). The cohesin complexes, encompassing the duplicated sister chromatids, are thought to build the basis of the axis in plants as well as in most, if not all, of other species. On top of cohesin complexes, at least three additional proteins are loaded, including the HORMA domain-containing protein (HORMAD) ASY1/PAIR2 (homolog of Hop1 in yeast, HORMAD1/2 in mammals), and two coiled-coil proteins known as the ‘axis core’ that affect ASY1 localization, i.e., the linker protein ASY3/PAIR3 (homolog of Red1 in yeast, SYCP2 in mammals) and ASY4 (homolog of SYCP3 in mammals) (Armstrong, 2002; Wojtasz *et al*, 2009; Hollingsworth & Johnson, 1993; Fukuda *et al*, 2010; Lammers *et al*, 1994; Ferdous *et al*, 2012; Chambon *et al*, 2018). The chromosome axis tethers and organizes the sister chromatids to form a higher-order structure of linear DNA loop-arrays and promotes efficient DSB formation, interhomolog-biased repair, and synaptonemal complex installation (Xue *et al*, 2019; Hollingsworth & Ponte, 1997; Niu *et al*, 2005; Goodyer *et al*, 2008; Carballo *et al*, 2008; Hunter, 2015; Lambing *et al*, 2020a, 2020b). Plants lacking any of these axis components show univalents in metaphase I meiocytes and/or altered CO distribution, highlighting the importance of chromosome axis for CO patterning (Lambing *et al*, 2020b; Sanchez-Moran *et al*, 2007; Ferdous *et al*, 2012; Armstrong, 2002; Daniel *et al*, 2011; Dio *et al*, 2023; Boden *et al*, 2009; Cuacos *et al*, 2021; Chambon *et al*, 2018; Pochon *et al*, 2022).

A crucial question concerning CO regulation is how CO assurance and interference are balanced. Recent studies show that loss of the transverse filament protein of the SC, ZYP1 in *Arabidopsis,* compromises CO assurance and abolishes interference, resulting in an ∼50% increase in class I COs, accompanied by ∼10 to 20% of metaphase I cells containing one pair of univalents (Capilla-Pérez *et al*, 2021; France *et al*, 2021; Yang *et al*, 2022). Nonetheless, the question to which extent this increase of class I COs is attributed to the loss of ZYP1 *per se* or to a defective remodeling of chromosome axis in general, remains obscure (Yang *et al*, 2022). Besides ZYP1, previous studies also provide insight into the role of ASY1 in CO assurance and interference (Lambing *et al*, 2020a; Pochon *et al*, 2022). In *Arabidopsis asy1* mutants which have a severely defective synapsis, CO interference is reported to be undetectable, and COs mainly locate and cluster at telomeric regions in contrast to more wildly spaced COs in the wildtype (Lambing *et al*, 2020a; Pochon *et al*, 2022).

Here, we carry out a fine-grained dissection for the function of chromosome axis protein ASY3 in CO formation through making use of the tetraploid nature of *Brassica napus* which harbors two copies of *ASY3* and thus four alleles offering the possibility to address the dosage dependency of ASY3 and other regulators. Cytological analyses in a series of *asy3* mutants where different number of *ASY3* alleles are mutated, show that the simultaneous mutation of all four *ASY3* alleles results in defective HEI10 loading and synapsis, and drastic reduction of chiasmata, which is largely rescued by the presence of only one functional *ASY3* allele, supporting the key role of ASY3 in implementing COs (Ferdous *et al*, 2012). Strikingly, while the number of class I COs in mutants with two functional *ASY3* alleles is comparable to that in the wildtype, this number is significantly increased in mutants with one functional *ASY3* allele, indicating that reducing ASY3 dosage increases CO formation. Moreover, the class I COs on each bivalent in mutants with one functional *ASY3* allele follows a Poisson-type distribution, indicating a strong attenuation of CO interference. Our results demonstrate the dosage-dependent distinct effects of ASY3 on CO formation and provide insights into the function and mechanism of chromosome axis/SC in CO patterning.

## Results

### Generation of *asy3* mutants in *Brassica napus*

By performing BLAST analysis using the *Brassica napus* multi-omics information resource database (https://yanglab.hzau.edu.cn/BnIR) (Yang *et al*, 2023; Song *et al*, 2020), two homologs (termed BnaASY3) of the *Arabidopsis* ASY3 protein were identified in *Brassica napus*. Each A and C sub-genomes contain one homolog: referred to as *BnaA05.ASY3* (*BnaA05g00870D*) located at chromosome A05 and *BnaC04.ASY3* (*BnaC04g00500D*) at chromosome C04. *BnaA05.ASY3* and *BnaC04.ASY3* exhibit a protein identity of 93.74% between each other, which share 75.13% and 75.16% protein identity with AtASY3, respectively (Supplemental Figure 1A).

To explore the function of ASY3, we applied the CRISPR-Cas9 based gene editing approach in the spring-type cultivar *Westar*, and identified from the 30 T0 generation of transformants two independent insertion mutant lines where monoallelic mutations are identified in both *BnaA05.ASY3* and *BnaC04.ASY3* genes (called *Bnaasy3-1* and *Bnaasy3-2*) (Figure 1A). The insertions in both *Bnaasy3-1* and *Bnaasy3-2* mutation lines result in premature translational termination of two *ASY3* copies, producing likely only truncated versions that contain only amino acids of the N-terminal regions (Supplemental Figure 1, red asterisks). For each line, we obtained different genotypes from a T1 segregating population, i.e., the mutants with four alleles of *ASY3* mutated (called *asy3-1^aacc^* and *asy3-2^aacc^*), mutants having only the two *ASY3* alleles of the A sub-genome mutated (called called *asy3-1^aaCC^* and *asy3-2^aaCC^*), and mutants having the two *ASY3* alleles of the A sub-genome plus one one allele of the C sub-genome mutated (named *asy3-1^aaCc^* and *asy3-2^aaCc^*). Unfortunately, for all T0 transformants, the two *ASY3* alleles of the A sub-genome harbor either no mutation or homozygous mutation (monoallelic or biallelic mutations), we, therefore, did not obtain such mutants of *asy3^AAcc^* and *asy3^Aacc^*.

**Figure 1.**
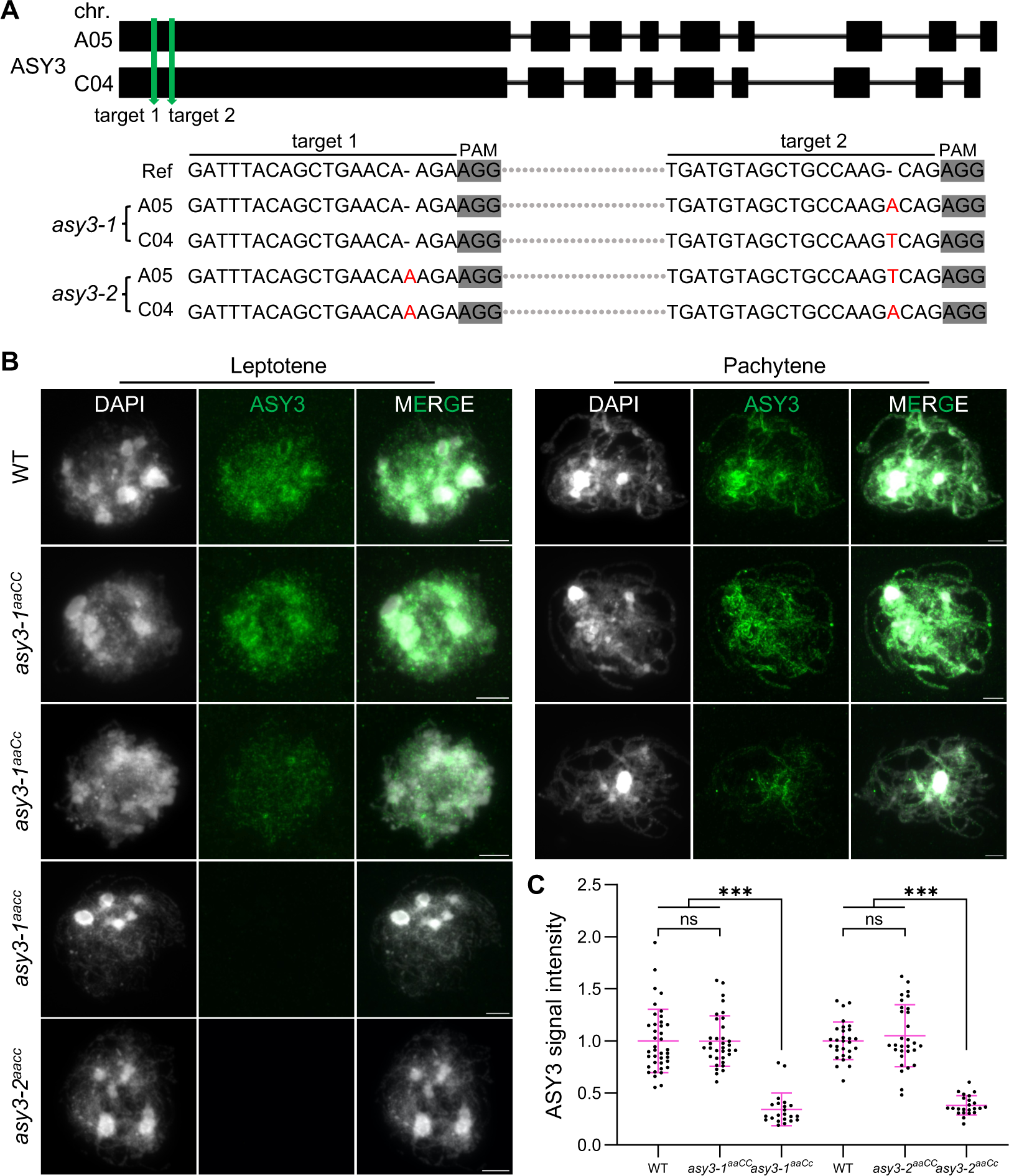
Generation of *asy3* null and hypomorphic mutants in *Brassica napus*. (A) Generation and identification of two *asy3* mutant lines by CRISPR-Cas9. (B) Immunostaining of ASY3 in male meiocytes of WT, *asy3-1^aaCC^*, *asy3-1^aaCc^*, *asy3-1^aacc^*, and *asy3-2^aacc^*mutant plants at leptotene and pachytene (or -like). Bars: 5µm. (C) Relative ASY3 signal intensity in WT, *asy3-1^aaCC^*, *asy3-1^aaCc^*, *asy3-2^aaCC^*, and *asy3-2^aaCc^* mutant plants at leptotene. The comparisons of signal intensity of WT, *asy3-1^aaCC^*, and *asy3-1^aaCc^* and WT, *asy3-2^aaCC^*, and *asy3-2^aaCc^* mutant plants were plotted independently.Asterisks indicate significant difference (Game-Howell’s multiple comparisons test, *p*<0.001).

To study the level of ASY3 accumulation in these different *asy3* mutants, we performed quantitative immunostaining analyses with an ASY3 antibody in male meiocytes (see method). The absence of the signal of ASY3 in *asy3-1^aacc^* and *asy3-2^aacc^* mutants validates the specificity of the antibody and confirms the complete loss of ASY3 function (Figure 1B). In wildtype, ASY3 accumulates along the chromosome axis at leptotene and at pachytene, two ASY3-labeled axes of the homologous chromosomes co-align and synapse resulting in thicker threads clearly visible in the immunostainings (Figure 1B). A similar ASY3 localization pattern was detected in *asy3-1^aaCC^* and *asy3-2^aaCC^* mutant alleles (Figure 1B, Supplemental figure 2A). In *asy3-1^aaCc^* and *asy3-2^aaCc^* mutants, where only one *ASY3* allele is functional, the thread-like structures of ASY3 were still observed along chromosomes at leptotene. However, the signal intensity of ASY3 in *asy3-1^aaCc^* and *asy3-2^aaCc^* was clearly weakened compared to the wildtype and *asy3^aaCC^* (Figure 1B, Supplemental Figure 2A). This reduction of the signal intensity of ASY3 in *asy3^aaCc^*persists to pachytene (or -like) cells where the ASY3 signal shows a patchier pattern compared to the wildtype and *asy3^aaCC^* plants (Figure 1B, Supplemental Figure 2A). We quantified the signal intensity of ASY3 during leptotene and pachytene (or -like) and observed a significant reduction of ASY3 dosage in *asy3-1^aaCc^* (∼ 65.77% decrease at leptotene, ∼ 61.71% at pachytene) and *asy3-2^aaCc^* (∼ 61.99% decrease at leptotene) compared to the wildtype (Game-Howell’s multiple comparisons test, *p*<0.001), while no significant difference was found between genotypes of *asy3^aaCC^* and wildtype at both leptotene and pachytene nuclei (Figure 1C, Supplemental Figure 2B).

### ASY3 dosage-dependent effects on the chromosome localization of ASY1

*Arabidopsis* ASY3 and its orthologs in other organisms, e.g., yeast, mouse, and rice, are required for proper chromosome recruitment and localization of the HORMAD protein ASY1 through a physical interaction of the N-terminal domain of ASY3 (known as closure motif, indicated in Supplemental figure 1A) and the HORMA domain of ASY1 (Yang *et al*, 2020b, 2020a; West *et al*, 2018; Rosenberg & Corbett, 2015; West *et al*, 2019; Kolas *et al*, 2004; Wang *et al*, 2011). We confirmed that this interaction of BnaASY3 closure motif (1-32 aa) to BnaASY1 HORMA domain (1-300 aa) is conserved in *Brassica napus* by using the yeast two-hybrid assay (Supplemental Figure 1B), supporting the potential role of ASY3 in ASY1 localization.

Next, to investigate the detailed effects of BnaASY3 on ASY1 localization, we performed an immunostaining analysis of male meiocytes using an antibody against ASY1. We verified the specificity of the ASY1 antibody using *Arabidopsis asy1-4* mutants (Supplemental figure 3A). In wildtype of *Brassica napus*, ASY1 forms linear structures along the chromosomes at leptotene and follows largely the REC8-labelled axes (Figure 2).

**Figure 2.**
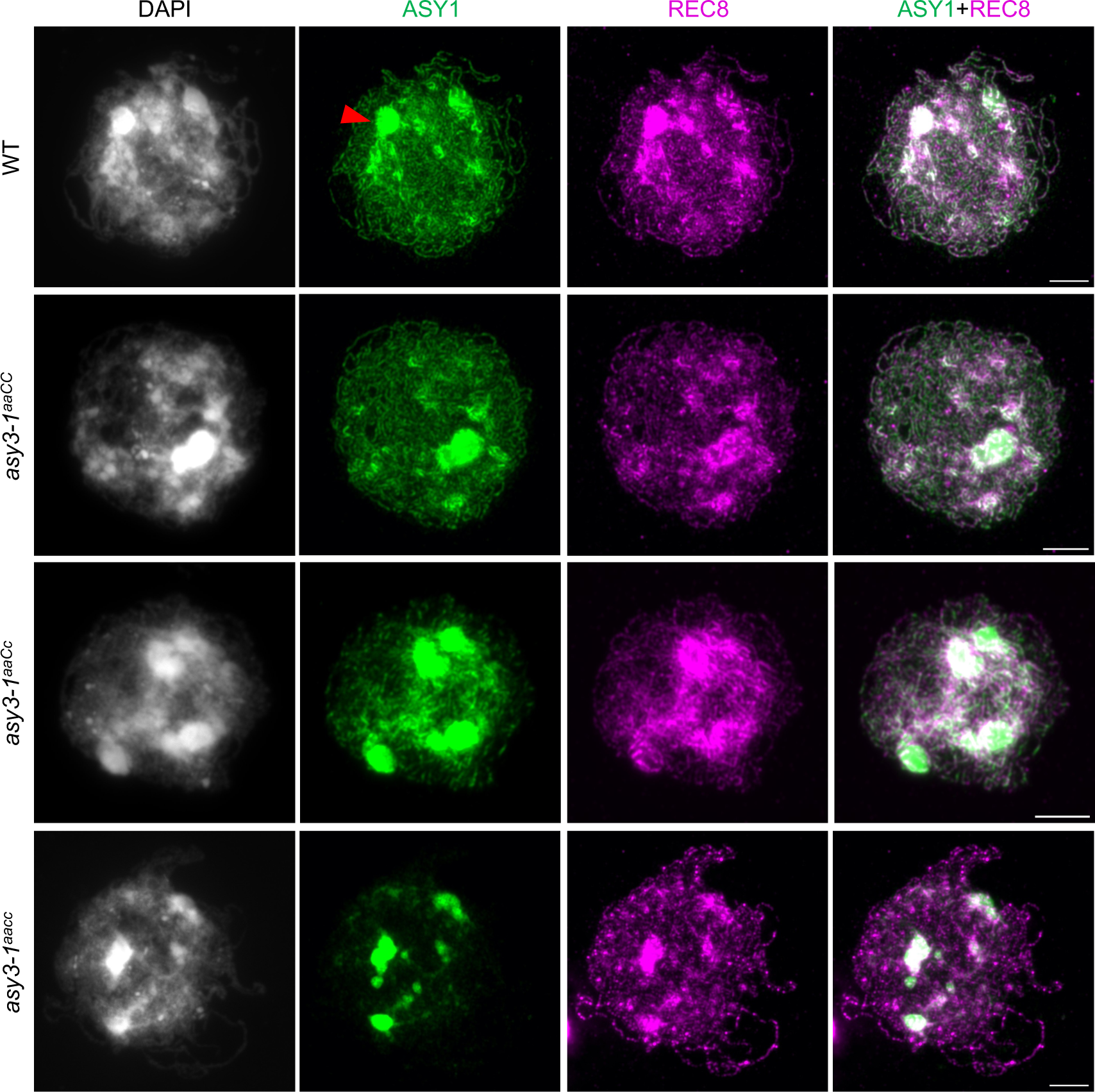
Immunolocalization of ASY1 and REC8 in male meiocytes of wildtype, *asy3-1^aaCC^*, *asy3-1^aaCc^*, and *asy3-1^aacc^*mutant plants at early prophase I (leptotene or leptotene-like). Red arrow indicates the “blob”-like signal that was overexposed and thus removed from the quantification of signal intensity shown in supplemental figure 4B. Bars: 5µm.

In *asy3^aacc^*, REC8 localization is normal whereas ASY1 is strongly diminished on most of the chromosome regions, consistent with an important role of ASY3 for proper recruitment and/or extension of ASY1 on chromosomes. Notably, a weak signal of ASY1 was still detected on chromosomes, showing a pattern of dotty signal or non-linear structures of short stretches. This observation is consistent with previous report in *Arabidopsis asy3* mutant, indicating that ASY1 may be able to bind to DNA independently of ASY3, as its homolog HOP1 in yeast (Khan *et al*, 2013; Kironmai *et al*, 1998).

We further investigated the localization of ASY1 in genotypes of *asy3^aaCC^*and *asy3^aaCc^* plants. We found that in *asy3-1^aaCC^* and *asy3-2^aaCC^*, ASY1 is indistinguishably loaded in comparison to the wildtype (Figure 2, Supplemental figure 4A). This result is consistent with the quantification of ASY3 dosage in *asy3^aaCC^* (Figure 1C). However, in *asy3-1^aaCc^*and *asy3-2^aaCc^* mutants where ASY3 dosage is less than 40% of that in wildtype, ASY1 shows a patchier and less continuous pattern with reduced signal intensity (Figure 2; Supplemental figure 4A). The quantification shows that while the protein level of ASY1 is not altered in in *asy3-1^aaCC^* and *asy3-2^aaCC^*, it is reduced to 58.92% and 54.58% of the wild-type level in *asy3-1^aaCc^*and *asy3-2^aaCc^*, respectively (Supplemental figure 4B).

### ASY3 exhibits a dosage-dependent impact on meiosis and plant fertility

We next analyzed the dosage-dependent effect of ASY3 on plant fertility. As expected, the absence of ASY3 does not affect plant growth and development until flowering, consistent with a meiosis-specific role of ASY3 (Supplemental Figure 5A) (Yuan *et al*, 2009; Wang *et al*, 2011; Ferdous *et al*, 2012). At reproductive stage, the silique length becomes very short in *asy3^aacc^* mutants and only very few viable seeds per silique are produced in comparison to the wildtype (average 1.83 in *asy3-1^aacc^* and 1.37 in *asy3-2^aacc^* vs 24.98 in wildtype) (Figure 3A and B). This reduced silique length is partially rescued by the presence of one functional *ASY3* allele in *asy3^aaCc^* genotypes. Accordingly, the seed-set is also significantly increased in *asy3^aaCc^* compared to *asy3^aacc^*mutants, but remaining lower than that in wildtype (average 17.41 in *asy3-1^aaCc^* and 15.26 in *asy3-2^aaCc^* vs 24.98 in wildtype, *p*<0.001, Game-Howell’s multiple comparisons test) (Figure 3A and B). In *asy3^aaCC^*mutants, the silique length and seed number per silique are comparable to that in the wildtype (Figure 3A and B).

**Figure 3.**
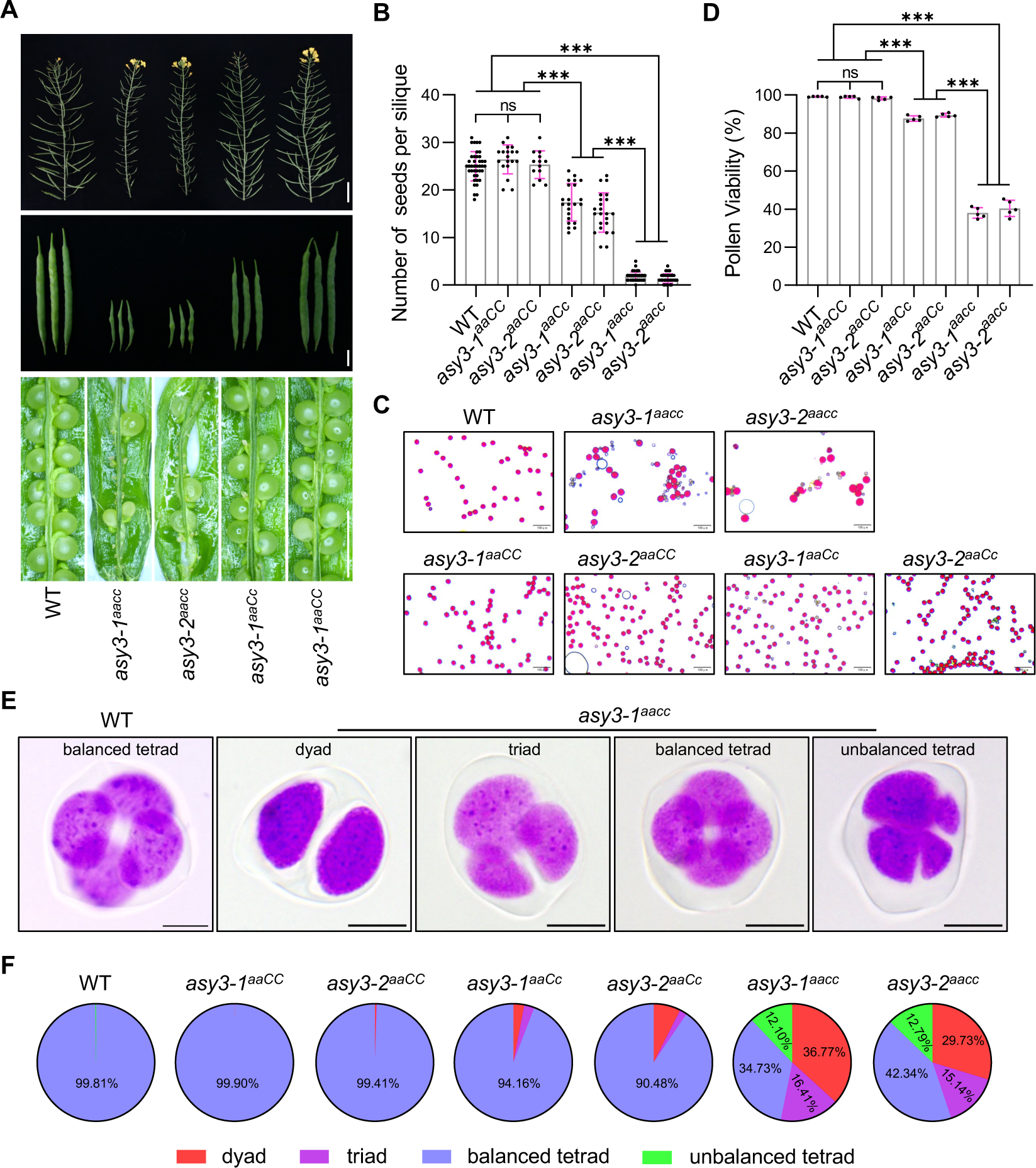
Phenotypic analysis of *asy3* mutants. (A) Main branches, siliques, and seed-sets in WT, *asy3-1^aacc^*, *asy3-2^aacc^*, *asy3-1^aaCc^*, and *asy3-1^aaCC^* mutant plants. Bars from top to bottom panels are 5 cm, 1 cm, and 5 mm, respectively. (B) Statistical analysis for the number of seeds per silique in WT, *asy3-1^aacc^*, *asy3-2^aacc^*, *asy3-1^aaCC^*, *asy3-2^aaCC^*, *asy3-1^aaCc^*, and *asy3-2^aaCc^* mutant plants. At least thirteen siliques from different plants were dissected and counted for each genotype. (C) Pollen staining of WT, *asy3-1^aacc^*, *asy3-2^aacc^*, *asy3-1^aaCC^*, *asy3-2^aaCC^*, *asy3-1^aaCc^*, and *asy3-2^aaCc^* mutant plants. Red staining indicates viable pollen and blue indicates dead pollen. Bars: 100µm (D) Pollen viability of WT, *asy3-1^aacc^*, *asy3-2^aacc^*, *asy3-1^aaCC^*, *asy3-2^aaCC^*, *asy3-1^aaCc^*, and *asy3-2^aaCc^* mutant plants. At least 4000 pollen grains were counted from different flowers for each genotype. (E) Examples of the staining of male meiotic products in WT and *asy3-1^aacc^* mutant plants. Bars: 5µm. (F) Pie charts showing the proportion of balanced tetrad, unbalanced tetrad, triad, and dyad in WT, *asy3-1^aacc^*, *asy3-2^aacc^*, *asy3-1^aaCC^*, *asy3-2^aaCC^*, *asy3-1^aaCc^*, and *asy3-2^aaCc^* mutant plants. Asterisks in (B and D) indicate significant difference (Game-Howell’s multiple comparisons test, *p*<0.001).

Staining pollen with Peterson solution revealed a similar ASY3 dosage-dependent impact on pollen viability (average 99.15% in WT vs 98.85% in *asy3-1^aaCC^*, 98.25% in *asy3-2^aaCC^*, 87.63% in *asy3-1^aaCc^*, 89.36% in *asy3-2^aaCc^*, 37.98% in *asy3-1^aacc^*, and 40.40% in *asy3-2^aacc^*) (Figure 3C and D). Subsequent tetrad analysis in male meiosis showed a severe disruption of meiotic products in *asy3^aacc^*mutants (65.27% abnormal tetrad in *asy3-1^aacc^*, and 57.66% in *asy3-2^aacc^* vs 0.19% in wildtype) (Figure 3E and F, Supplemental figure 5B). This defect in tetrad formation was largely reversed by the presence of one functional *ASY3* allele, producing only a small portion of defective tetrads compared to wildtype (5.84% in *asy3-1^aaCc^*, and 9.52% in *asy3-2^aaCc^* vs 0.19% in wildtype). In *asy3-1^aaCC^*and *asy3-2^aaCC^* mutants, the tetrad formation is indistinguishable from that in wildtype (Figure 3E and F, Supplemental figure 5B).

### Chromosome synapsis is sensitive to ASY3 dosage

For a detailed analysis of meiosis, we spread the chromosomes of male meiocytes. In the wildtype, the DAPI-stained chromatin exhibits a thin filament-like structure at leptotene and starts to pair with their homologs during zygotene (Figure 4A). As cells progress to pachytene, homologous chromosomes are co-aligned and synapsed (n=36 cells). In *asy3^aacc^* mutants, the chromosomes also show a thread-like structure at leptotene (Figure 4A, Supplemental figure 6A). However, similar to *Arabidopsis* and rice *asy3/pair3* mutants (Yuan *et al*, 2009; Wang *et al*, 2011; Ferdous *et al*, 2012), normal pachytene nuclei were never observed, and homologous chromosomes remain largely not co-aligned (n=50 cells in *asy3-1^aacc^*, and n=31 in *asy3-2^aacc^*) (Figure 4A, Supplemental figure 6A).

**Figure 4.**
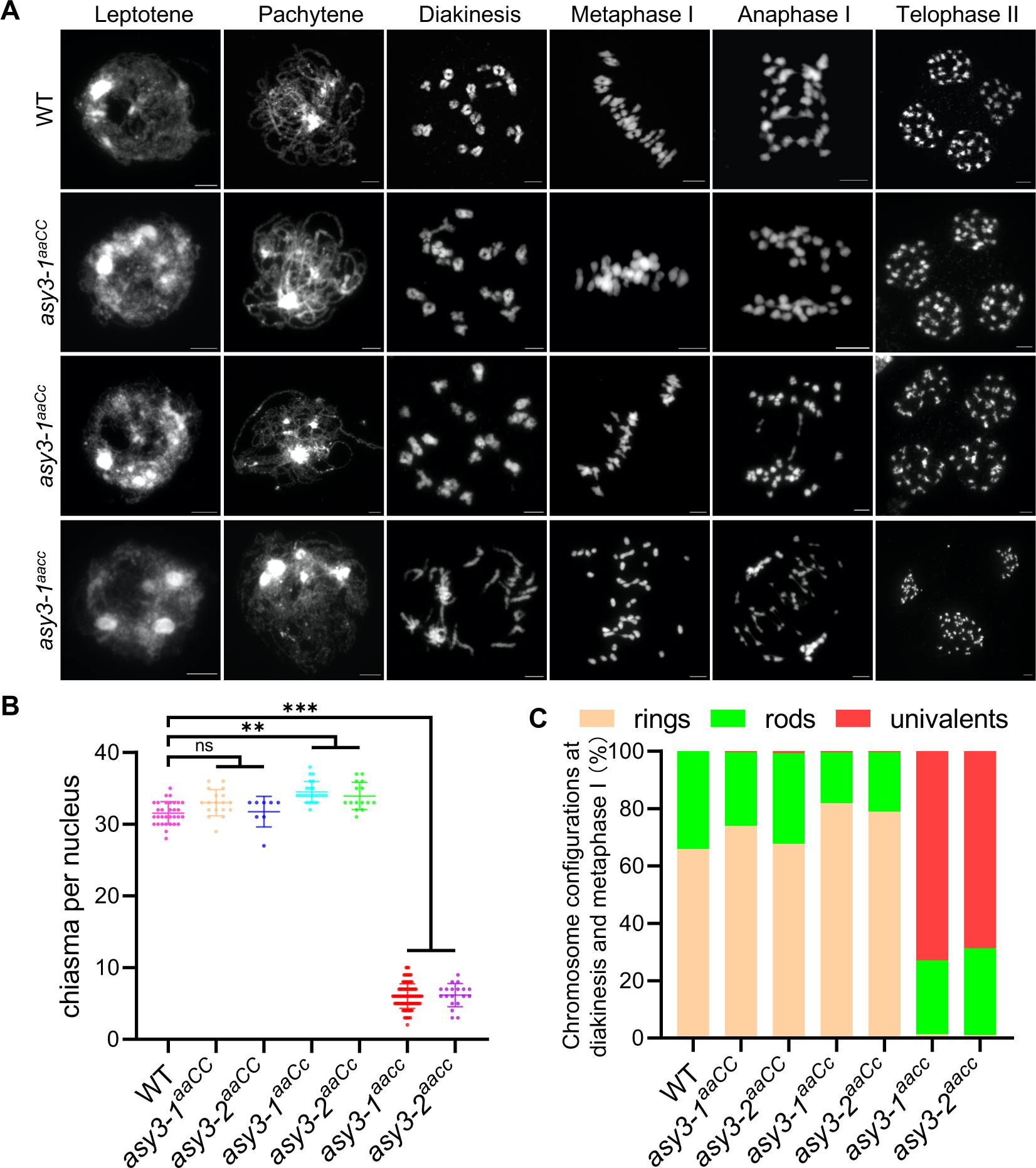
Analysis of meiotic chromosome behavior in wildtype and *asy3* mutants. (A) Chromosome spread analysis of male meiosis in WT, *asy3-1^aaCC^*, *asy3-1^aaCc^*, and *asy3-1^aacc^* mutant plants throughout meiosis. Bars: 5µm. (B) Scatter dot plot of the estimation of minimum chiasma number in WT, *asy3-1^aaCC^*, *asy3-2^aaCC^*, *asy3-1^aaCc^*, *asy3-2^aaCc^*, *asy3-1^aacc^*, and *asy3-2^aacc^* mutant plants. The statistical analysis was performed by one-way ANOVA with Games-Howell’s multiple comparisons test. **, *p*<0.01 and ***, *p*<0.001. (C) Distribution of chromosome configurations at diakinesis/metaphase I (ring, rod, or univalent) in WT, *asy3-1^aacc^*, *asy3-2^aacc^*, *asy3-1^aaCC^*, *asy3-2^aaCC^*, *asy3-1^aaCc^*, and *asy3-2^aaCc^*mutant plants.

In genotypes of *asy3^aaCC^*, chromosome juxtaposition at pachytene seems to be properly achieved in a large proportion of cells (79.31%, n=58 cells in *asy3-1^aaCC^*, and 80.70%, n=57 cells in *asy3-2^aaCC^*) (Figure 4A, Supplemental figure 6A), with the remaining cells showing only small regions of unpaired stretches (red arrows in Supplemental figure 6B). In *asy3^aaCc^* mutants, defects in chromosome coalignment and synapsis were more frequently observed and all pachytene cells showed some unpaired single chromosome threads while most of the chromosome regions were able to synapse (n=48 cells in *asy3-1^aaCc^*, and n=51 cells in *asy3-2^aaCc^*) (Figure 4A, Supplemental figure 6A). Noting that the ratio of cells with properly/fully co-aligned chromosomes could be a bit overestimated due to the low resolution of the DAPI-stained chromosomes.

To complement the above analysis, we performed co-immunolocalization of ZYP1 and REC8 in male meiosis at pachytene. In the wildtype, ZYP1 is assembled along the entire chromosome length, which links the REC8 axes (n=14) (Figure 5A). Corresponding to what we observed in chromosome spreads stained with DAPI, a large portion of cells of the *asy3^aaCC^* showed complete ZYP1 assembly (60.61% in *asy3-1^aaCC^*, n=32 cells and 61.90% in *asy3-2^aaCC^*, n=21) (Figure 5A, Supplemental figure 7A), while the rest of the nuclei harbor only short non-ZYP1 stained chromosome regions (arrowheads in Supplemental figure 7B). In *asy3^aaCc^* mutants, compared to the REC8-labeled axes, ZYP1 could be installed onto most regions of the chromosomes, although a complete SC assembly was never observed (n=20 cells in *asy3-1^aaCc^*, and n=21 cells in *asy3-2^aaCc^*) (arrowheads in Figure 5A, Supplemental figure 7A). These results indicate that the presence of only one functional allele of *ASY3* in *Brassica napus* largely recovers the pairing and synapsis defects induced by the absence of ASY3.

**Figure 5.**
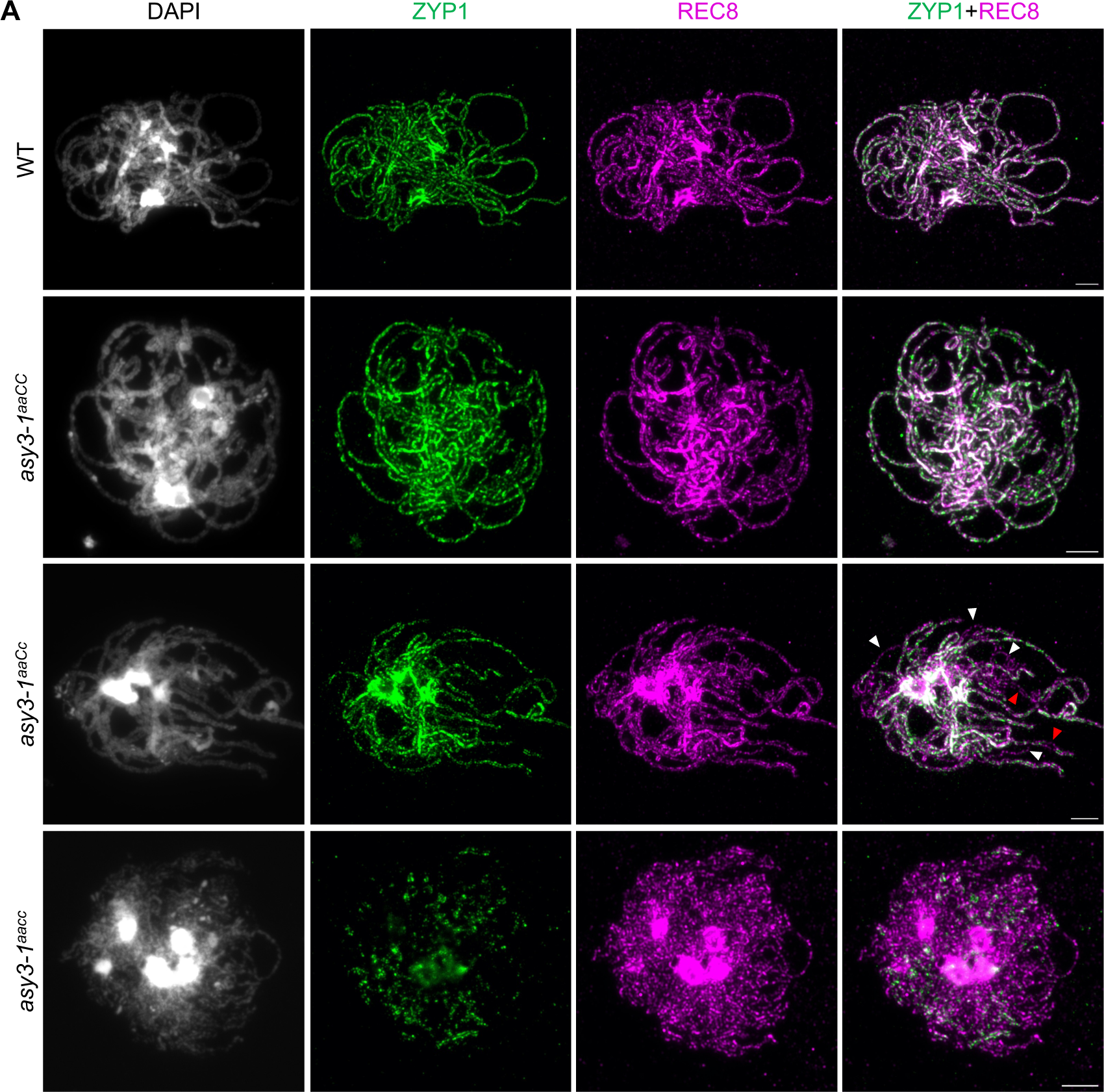
ASY3 dosage-dependent effects on chromosome synapsis. (A) Co-immunolocalization of ZYP1 and REC8 at pachytene in male meiocytes of WT, *asy3-1^aaCC^*, *asy3-1^aaCc^*, and *asy3-1^aacc^* mutant plants. White and red arrowheads indicate the single threads or coaligned regions that both have no ZYP1 signal, respectively. Bars: 5µm.

### ASY3 shows dosage-dependent different effects on chiasma formation

To ensure balanced chromosome segregation, each pair of homologs needs to have at least one CO, visible as a chiasma in chromosome spreads. In wildtype, in total 19 condensed bivalents were counted in all observed meiocytes of *Brassica napus* at diakinesis (n=48 cells), which were aligned at the equatorial plate at metaphase I to achieve balanced segregation (n=10 cells) (Figure 4A). However, in *asy3^aacc^* null mutants, a large amount of univalents were visible in all nuclei at diakinesis (average 27.71±2.80 univalents, n=49 cells in *asy3-1^aacc^* and 26.11±3.46 univalents, n=19 cells in *asy3-2^aacc^*) and could not move properly to the metaphase I plate in all observed cells (n=24 cells in *asy3-1^aacc^*, and n=20 cells in *asy3-2^aacc^*) (Figure 4A, Supplemental figure 6A). Consequently, chromosome segregation is unbalanced at anaphase I, accompanied by the appearance of chromosome bridges, presumably reflecting premature separation of sister chromatids (Figure 4A, Supplemental figure 6A). These results support the conclusion that ASY3 is needed for CO formation (Ferdous *et al*, 2012; Wang *et al*, 2011; Yuan *et al*, 2009).

Compared to *asy3^aacc^* mutants, the bivalent formation is largely achieved in both genotypes of *asy3^aaCC^* and *asy3^aaCc^* plants (Figure 4A, Supplemental figure 6A, Supplemental table 1). Occasionally, two univalent were observed, i.e., one pair of chromosomes failed to form a CO (7.5% in *asy3-1^aaCC^*, n=40 cells; 6.67% in *asy3-2^aaCC^*, n=15; 15.63% in *asy3-1^aaCc^*, n=64; and 14.29% in *asy3-2^aaCc^*, n=56) (arrowheads in Supplemental figure 6C). However, more than two univalents were never detected in *asy3^aaCC^* and *asy3^aaCc^* plants.

Next, we quantified the chiasma number based on the chromosome configurations at diakinesis/metaphase I: ring-type bivalent has at least two chiasmata while rod-type bivalent has one chiasma (Figure 4B, Supplemental table 1). Notably, this estimated number of chiasmata is possibly lower than the actual number of COs, especially in meiotic mutants. In wildtype, in average 31.53±1.56 chiasmata (n=32 cells) are formed, which contains 65.95% ring-type and 34.05% rod-type bivalents (Figure 4B and C) (n=32 cells). In *asy3^aacc^* mutants, 72.93% (n=49 cells) and 68.70% (n=19 cells) of chromosome pairs occurred as univalents in *asy3-1^aacc^* (6.04±1.71 chiasmata, n=100 cells) and *asy3-2^aacc^* (6.16±1.56 chiasmata, n=19 cells) mutants, respectively, with the remaining bivalents present mainly as the rod type (Figure 4A-C, Supplemental figure 6A). This defect of chiasma formation was almost fully recovered in *asy3^aaCC^* mutants in which the number of chiasmata reached wild-type level (33.00±1.76 chiasmata in *asy3-1^aaCC^*, n=18 cells and 31.75±1.98 chiasmata in *asy3-2^aaCC^*, n=8 vs 31.53±1.56 in wildtype, n=32) (Figure 4A and C, Supplemental figure 6A), despite a fraction of cells harbors one pair of univalents (7.5% in *asy3-1^aaCC^*, n=40 cells; 6.67% in *asy3-2^aaCC^*, n=15) (see above, arrowheads in Supplemental figure 6C).

Unexpectedly, we found that the ratio of ring-type bivalents in *asy3^aaCc^* mutants is dramatically increased (81.92 % in *asy3-1^aaCc^*, n=23 and 78.95% in *asy3-2^aaCc^*, n=15 vs 65.95% in wildtype, n=32), indicating an significant increase of total number of chiasmata (34.48±1.44 in *asy3-1^aaCc^*, n=23 and 33.93±1.84 in *asy3-2^aaCc^*, n=15 vs 31.53±1.56 in wildtype, n=32, Game-Howell’s multiple comparisons test, *p*<0.01) (Figure 4B and C, Supplemental table 1).

### ASY3 is required for the wild-type level of initial loading of HEI10

To further understand the molecular effect of a reduction of ASY3 dosage on meiotic recombination, we performed immunolocalization analysis for HEI10 at different stages of male meiosis (Supplemental table 2). HEI10 (also known as ZHP-3/4 in *C. elegans*), a ring type E3 ubiquitin ligase, belongs to the ZMM group of proteins. HEI10 displays a dynamic localization pattern during meiotic prophase I which seems conserved in many species including *Brassica napus*, i.e., it initially forms numerous small foci at zygotene and early pachytene that are progressively consolidated into large foci colocalizing with the class I CO sites at late pachytene, diplotene, and diakinesis (Pinzón *et al*, 2021; Mercier *et al*, 2015; Grandont *et al*, 2014; Chelysheva *et al*, 2012; Wang *et al*, 2012). The specificity of the HEI10 antibody is validated in the *Arabidopsis hei10-2* mutant (Supplemental figure 3B). Counting the number of HEI10 foci at zygotene and early pachytene, we found that the numbers of HEI10 foci in asy3*-1^aaCC^*(130.73±17.04, n=11), *asy3-2^aaCC^* (131.60±17.60, n=10), *asy3-1^aaCc^*(123.77±22.52, n=26), and *asy3-2^aaCc^* (126.81±31.38, n=21) mutants are not significantly different from that in wildtype (134.90±33.88, n=21) (Games-Howell’s multiple comparisons test, p>0.05) (Figure 6A and B, Supplemental figure 8A, Supplemental table 2), suggesting that the initial HEI10 loading is maintained at least when ASY3 protein amount is reduced to ∼30-40% of wildtype. However, this number is drastically reduced in *asy3-1^aacc^* (12.43±3.92, n=30) and *asy3-2^aacc^* (10.95±2.33, n=22) mutants (Figure 6A and B, Supplemental Figure 8A). Consistently, we saw that the initial HEI10 loading is also largely compromised in *Arabidopsis asy3-1* mutants (11.14±4.00 in *asy3-1*, n=21 vs 37.38±8.35 in wildtype, n=24) (student’s t-test, *p*<0.001) (Supplemental figure 8B and C). These results suggest that ASY3 is important for the initial recruitment of HEI10 onto chromosomes at early prophase I in both *Brassica napus* and *Arabidopsis*.

**Figure 6.**
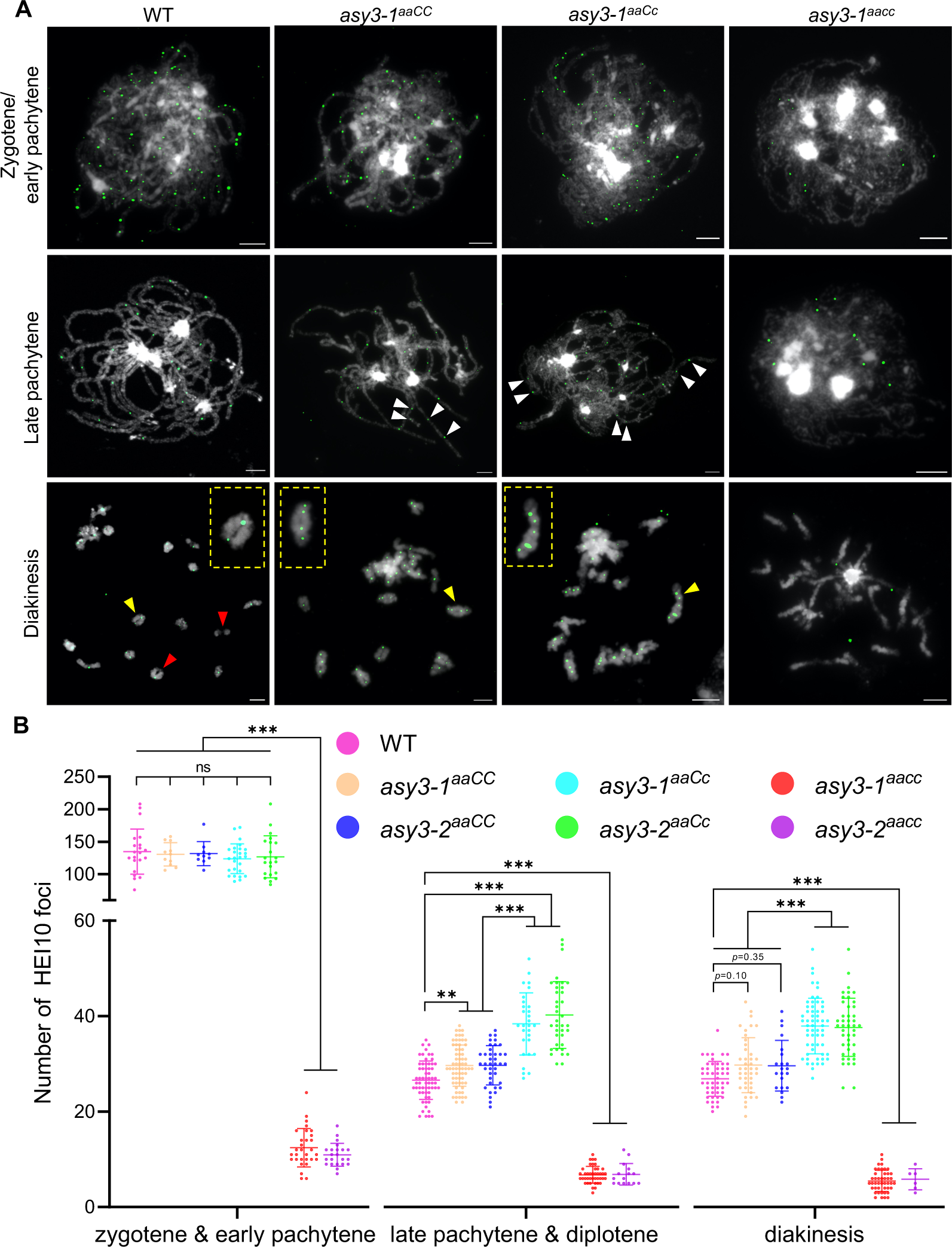
HEI10 localization in wildtype and *asy3* mutants. (A) Immunostaining of HEI10 at early prophase I (zygotene/early pachytene), late pachytene, and diakinesis in male meiocytes of WT, *asy3-1^aaCC^*, *asy3-1^aaCc^*, and *asy3-1^aacc^* mutant plants. White arrowheads in the images of late pachytene indicate the closely localized HEI10 foci along one paired chromosome. Red arrowheads indicate the position of class II COs. Yellow arrowheads depict the magnified bivalents shown in the yellow rectangles. Bars: 5µm. (B) Quantification of the number of HEI10 foci at zygotene/pachytene, late pachytene/diplotene, and diakinesis in male meiocytes of WT, *asy3-1^aaCC^*, *asy3-2^aaCC^*, *asy3-1^aaCc^*, *asy3-2^aaCc^*, *asy3-1^aacc^*, and *asy3-2^aacc^*mutant plants. Games-Howell’s multiple comparisons test, ** *p*<0.01, and *** *p*<0.001.

### ASY3 dosage modulates the formation of class I crossovers

Similar to *Arabidopsis*, at late pachytene and diplotene, the number of HEI10 foci in *asy3-1^aacc^* (6.76±1.75, n=41 cells) and *asy3-2^aacc^* (6.87±2.19, n=15) is strongly reduced compared to that of wildtype (26.59±3.99, n=61); at diakinesis the number of HEI10 foci is decreased from 26.88±3.62 (n=48) in the wildtype to 5.48±2.20 (n=44) in *asy3-1^aacc^* and 5.83±2.03 (n=6) in *asy3-2^aacc^* (Games-Howell’s multiple comparisons test, *p*<0.001) (Figure 6A and B, Supplemental Figure 8A, Supplemental table 2). This suggests that ASY3 is required for the formation of interference-sensitive COs, consolidating its role in promoting synapsis and chiasma formation.

Unexpectedly, in *asy3^aaCC^*mutant alleles, the number of HEI10 foci is slightly, yet significantly increased at late pachytene and diplotene (29.65±4.30 in *asy3-1^aaCC^*, n=66 and 29.72±4.06 in *asy3-2^aaCC^*, n=39 vs 26.59±3.99 in wildtype, n=61, Games-Howell’s multiple comparisons test, *p*<0.01); at diakinesis HEI10 foci still show a slight increase though not statistically significant in both asy3*-1^aaCC^* (29.76±5.69, n=41 vs 26.88±3.62 in wildtype, n=48, *p*=0.10) and *asy3-2^aaCC^* (29.62±5.16, n=21, *p*=0.35) (Games-Howell’s multiple comparisons test) (Figure 6A and B, Supplemental Figure 8A).

More strikingly, we found that the number of HEI10 foci in *asy3-1^aaCc^* (38.38±6.40, n=29) and *asy3-2^aaCc^*(40.24±6.91, n=37) are even more dramatically elevated at late pachytene/diplotene, i.e., 44% and 51% increase in comparison to wildtype, respectively (Figure 6A and B, Supplemental Figure 8A). This significant elevation of HEI10 foci in *asy3-1^aaCc^* and *asy3-2^aaCc^* is further confirmed in meiocytes at diakinesis (37.91±5.78 in *asy3-1^aacc^*, n=58 and 37.66±6.03 in *asy3-2^aacc^*, n=44 vs 26.88±3.62 in wildtype, n=48, 41% and 40% increase in comparison to wildtype, respectively) (Games-Howell’s multiple comparisons test, *p*<0.001) (Figure 6A and B, Supplemental Figure 8A). This result is consistent with the increase of chiasmata in *asy3^aaCc^* mutants (Figure 4C). Altogether, these results show that ASY3 modulates the formation of class I COs in a dosage-dependent manner.

### Interference-insensitive COs are reduced in *asy3^aacc^*, but not in *asy3^aaCC^* and *asy3^aaCc^*

To understand whether ASY3 also plays a role in the formation of class II (interference-insensitive) COs, we estimated the amount of class II COs by analyzing the configurations of bivalents labeled by HEI10 at diakinesis, i.e., non-HEI10 labeled chiasmata were treated as class II COs (e.g., red arrowheads in Figure 6A). In this way, the estimated total amount of COs equals the HEI10 foci on bivalents of diakinesis plus the amount of class II COs (i.e., non-HEI10 labeled chiasmata). Based on this estimation, the numbers of class II and total COs in wildtype were 6.55±3.09 (n=29) and 33.97±2.54 (n=29), respectively (Figure 7A and B). We found that the numbers of all COs and class II COs were both not significantly altered in *asy3^aaCC^* mutants (Figure 7A and B, Supplemental table 2). However, besides the drastic decrease of total COs (Figure 7A), the amount of class II COs was significantly reduced to ∼ 0.56 and ∼ 0.32 in *asy3-1^aacc^* and *asy3-2^aacc^*null mutants, respectively (Supplemental table 2), suggesting that ASY3 in *Brassica napus* is likely required for the formation of both the interference-sensitive and -insensitive COs, as in *Arabidopsis* (Ferdous *et al*, 2012).

**Figure 7.**
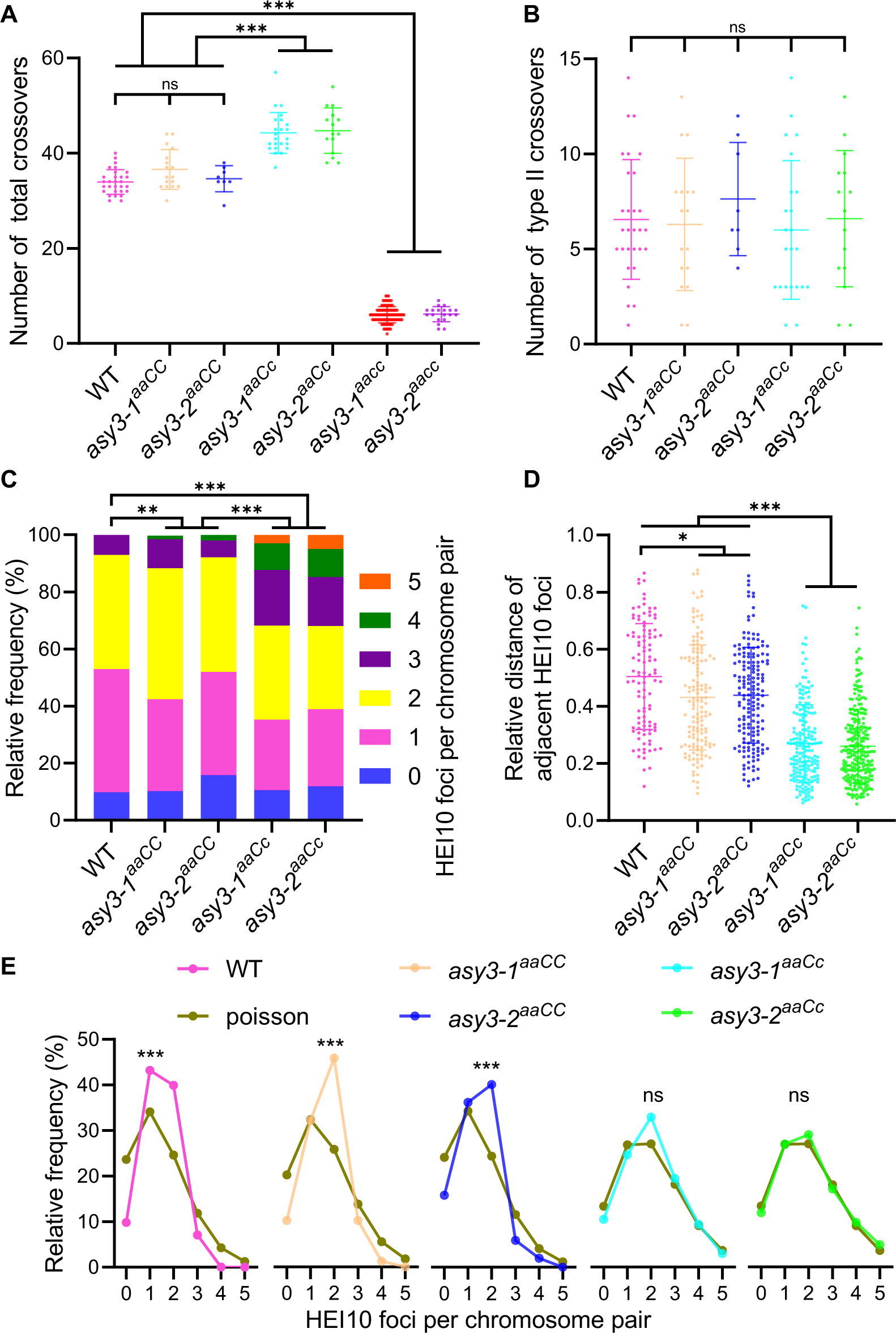
ASY3 dosage modulates crossover number and distribution. (A) Quantification of the estimated number of total COs in male meiocytes of WT, *asy3-1^aaCC^*, *asy3-2^aaCC^*, *asy3-1^aaCc^*, *asy3-2^aaCc^*, *asy3-1^aacc^*, and *asy3-2^aacc^* mutant plants. Games-Howell’s multiple comparisons test. (B) Quantification of the estimated number of Class II COs in male meiocytes of WT, *asy3-1^aaCC^*, *asy3-2^aaCC^*, *asy3-1^aaCc^*, and *asy3-2^aaCc^* mutant plants. Tukey’s multiple comparisons test. (C) Distribution of the number of HEI10 foci in male meiocytes of WT, *asy3-1^aaCC^*, *asy3-2^aaCC^*, *asy3-1^aaCc^*, and *asy3-2^aaCc^* mutant plants. Games-Howell’s multiple comparisons test. (D) Relative distance of adjacent HEI10 foci on each bivalent at diakinesis in male meiocytes of WT, *asy3-1^aaCC^*, *asy3-2^aaCC^*, *asy3-1^aaCc^*, and *asy3-2^aaCc^*mutant plants. (E) Comparison of the observed and Poisson-predicted distributions of the number of HEI10 foci per chromosome pair (bivalent) in male meiocytes of WT, *asy3-1^aaCC^*, *asy3-2^aaCC^*, *asy3-1^aaCc^*, and *asy3-2^aaCc^* mutant plants. Chi-square test. * *p*<0.05, ** *p*<0.01, and *** *p*<0.001.

Notably, we observed that, despite the marked increase of the class I COs in *asy3^aaCc^* (Figure 6B), the number of class II COs was not altered compared to wildtype (6.00±3.56 in *asy3-1^aaCc^*, n=23, *p*=0.98; 6.60±3.46 in *asy3-2^aaCc^*, n=15, *p*=0.99, Tukey’s multiple comparisons test) (Figure 7B), suggesting that the formation of class II COs is likely not sensitive to a moderate reduction of ASY3 dosage. Therefore, we propose that an appropriate modulation of ASY3 dosage could increase the formation of class I COs without affecting class II COs.

### ASY3 dosage modifies crossover interference

CO interference spaces adjacent COs and leads to the ubiquitous observation that each bivalent typically contains only 1-3 COs (Zickler & Kleckner, 2015; Mercier *et al*, 2015). In wild-type male meiocytes of *Brassica napus*, we observed typically only one large HEI10 focal point consolidated along a long stretch of synapsed chromosomes at late pachytene (Figure 6A), consistent with previous studies in rapeseed and other organisms (Grandont *et al*, 2014; Gonzalo *et al*, 2019; Desjardins *et al*, 2022; Morgan *et al*, 2021; Wang *et al*, 2012; Chelysheva *et al*, 2012). However, the phenomenon of two or even more HEI10 foci localized on a short interval of synapsed chromosomes at late pachytene was frequently observed in *asy3^aaCC^* (Figure 6A). This phenomenon became more obvious in *asy3^aaCc^* alleles where all nuclei (n=29 in *asy3-1^aaCc^*, and n=37 in *asy3-2^aaCc^*) showed more closely localized HEI10 foci along one synapsed/co-aligned chromosome pair compared to that in wildtype (white arrows in Figure 6A, Supplemental figure 8A). These results suggest that CO interference is likely less effective as ASY3 dosage decreases.

Therefore, we analyzed and plotted the CO pattern by counting the number of HEI10 foci per bivalent at diakinesis (Figure 6A and 7C, Supplemental figure 8A). In wildtype, the majority of bivalents contained one (43.19%) or two (39.93%) HEI10 foci (n=551 bivalents); the remaining bivalents had either three (7.08%) or no HEI10 foci (9.80%) (Figure 7C). In *asy3^aaCC^*, a small proportion of bivalents had 4 HEI10 foci (1.24%, n = 323 bivalents in *asy3-1^aaCC^*, 1.97%, n=152 bivalents in *asy3-2^aaCC^*), which was never observed in wildtype (n=551 bivalents) (Figure 7C). More strikingly, the ratio of bivalents with 3 or more COs (HEI10 foci) was dramatically increased in *asy3-1^aaCc^*(31.81%, n = 437 bivalents) and *asy3-2^aaCc^* (31.93%, n=285 bivalents) compared to wildtype (7.08%) (n=551 bivalents), and a portion of bivalents even contained five HEI10 foci (2.97% in *asy3-1^aaCc^* and 4.91% in *asy3-2^aaCc^*) (Figure 6A and 7C, Supplemental figure 8A).

Statistically, the number of HEI10 foci per bivalent in *asy3^aaCC^*was significantly distinct from that in wildtype (Chi-square test, *p=*0.001*7* and 0.0026, respectively) (Figure 7C). Notably, there was a strongly altered distribution of HEI10 focus number per chromosome pair in *asy3^aaCc^* compared to both wild-type and *asy3^aaCC^* plants (Chi-square test, *p<*0.001) (Figure 7C). Moreover, we measured the relative distance of adjacent HEI10 foci at diakinesis (see method). We found that compared to the wildtype (0.50±0.18, n=106), this distance became slightly yet significantly closer in *asy3^aaCC^* (0.43±0.18 in *asy3-1^aaCC^*, n=142; 0.44±0.17 in *asy3-2^aaCc^*, n=168, Games-Howell’s multiple comparisons test, *p*<0.05), and was drastically reduced in *asy3^aaCc^* (0.27±0.14 in *asy3-1^aaCc^*, n=188; 0.26±0.13 in *asy3-2^aaCc^*, n=251, Games-Howell’s multiple comparisons test, *p*<0.001) (Figure 7D), implying further the attenuation of CO interference.

Furthermore, we performed a distribution analysis for HEI10 focus number per bivalent at diakinesis. In wildtype, the distribution of HEI10 focus number per chromosome pair significantly deviated from an expected Poisson random distribution (chi-square test, wildtype vs Poisson, *p*=6.48E-31) (Figure 7E). In *asy3^aaCC^*, HEI10 distribution was also significantly distinct from the Poisson distribution (chi-square test, *asy3-1^aaCC^* vs Poisson, *p*=5.98E-17; *asy3-2^aaCC^* vs Poisson, *p* = 4.28E-5) (Figure 7E). Considering the shorter distance of inter-HEI10 foci and modified HEI10 distribution on diakinesis bivalents compared to wildtype (Figure 7C and D), we conclude that CO interference is mildly reduced in *asy3^aaCC^*. Strikingly, the observed HEI10 distribution per bivalent in *asy3^aaCc^*largely fits the expected values of Poisson distribution (chi-square test, *asy3-1^aaCc^* vs Poisson, *p*=0.08; *asy3-2^aaCc^*vs Poisson, *p*=0.78) (Figure 7E), suggesting that CO interference is compromised when ASY3 dosage is reduced to less than half of the wildtype level.

## Discussion

The chromosome axis plays a crucial role in CO patterning through providing a platform for recruiting the recombination machinery and transmitting the force of CO interference (Zhang *et al*, 2014b; Zickler & Kleckner, 2015; Lambing *et al*, 2020a, 2020b; Sanchez-Moran *et al*, 2007). Previous work in *Arabidopsis* has shown that in both *asy3* and *asy1* mutants, CO formation is severely compromised, resulting in a large reduction in bivalent formation (Ferdous *et al*, 2012; Armstrong, 2002; Dio *et al*, 2023; Cuacos *et al*, 2021). The remaining COs in *asy1* mutants have been shown to be clustered in telomere-proximal regions and exhibit compromised interference, suggesting a role of ASY1 in antagonizing telomere-led recombination and promoting crossover formation in interstitial chromosome arms (Lambing *et al*, 2020a; Pochon *et al*, 2022). In *Arabidopsis asy1/+* (∼21% reduction in ASY1 loading) and *asy3/+* (∼25% reduction in ASY1 loading) heterozygous mutants, the CO landscape is also remodeled, with a shift toward the distal subtelomeres, at the expense of interstitial and pericentromeric regions (Lambing *et al*, 2020a). However, the global CO numbers and interference are still maintained in *Arabidopsis asy1/+* and *asy3/+* heterozygotes with full pairing and synapsis occurring (Lambing *et al*, 2020a).

### ASY3 has dosage-dependent diverse effects on recombination

Here, we make use of the tetraploid nature of *Brassica napus* to generate an allelic series of *asy3* mutants allowing a fine-grained dissection of the dosage-function relationship, and address how the axis protein ASY3 modulates CO frequency and distribution (Figure 8). In wildtype, ASY3 promotes synapsis and ensures that each pair of chromosomes forms the obligatory CO. At the same time, a high/wild-type level of ASY3 facilitates CO interference. Consequently, each pair of chromosomes typically obtains only 1 to 3 COs (Mercier *et al*, 2015; Hunter, 2015). In *asy3^aacc^* mutants where ASY3 is completely absent, CO formation is severely disrupted due to the loss of CO promotion mediated by ASY3, indicating that ASY3 is crucial for CO generation.

**Figure 8.**
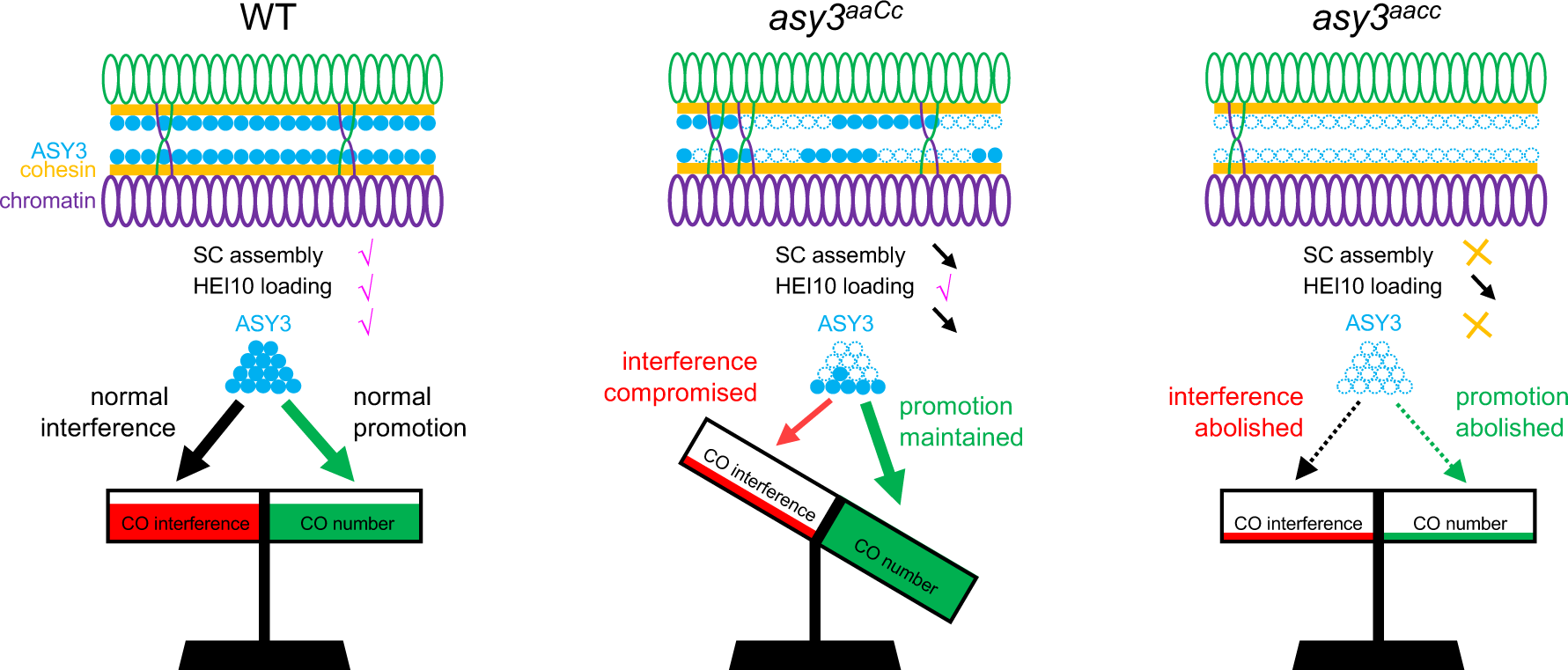
Model for the crossover formation with different ASY3 dosages. In wildtype, high level of ASY3 ensures the fidelity of HEI10 loading and SC assembly that, on the one hand, promotes the interhomolog recombination and, on the other hand, provides the platform for HEI10 dynamic coarsening, thus implementing both the CO promotion and interference. In the absence of ASY3, SC assembly is disrupted, and initial HEI10 loading is interfered, which largely compromises both the CO promotion and interference, resulting in the drastic decrease of COs. In *asy3^aaCc^* where ASY3 dosage is reduced to ∼30%-40% of the wild-type level, CO interference is compromised, whereas HEI10 loading and SC assembly are still maintained to a large extent, resulting in the global increase of COs.

Interestingly, our data show that when ASY3 dosage is reduced to ∼30 to 40% of the wildtype, as seen in the *asy3^aaCc^* plants, its function in promoting pairing and CO formation is largely maintained (Figure 4 and 5), but the competence to orchestrate interference is almost abolished (see below), thus leading to an increase of class I COs while only a mild defect in CO assurance occurs (Figure 8). This could indicate the existence of a threshold over which ASY3 imposes CO interference (directly or indirectly) and in turn, constraining the formation of excess COs.

### Mechanisms of ASY3 for CO formation

The chromosome axis has been shown to be important for meiotic recombination in many species as diverse as budding yeast, mice, *Drosophila*, *C. elegans*, *Arabidopsis*, rice, and wheat (Hollingsworth *et al*, 1990; Latypov *et al*, 2010; Hollingsworth & Johnson, 1993; Dubois *et al*, 2019; Wang *et al*, 2011; Wojtasz *et al*, 2009; Couteau & Zetka, 2005; Martinez-Perez & Villeneuve, 2005; Goodyer *et al*, 2008; Sanchez-Moran *et al*, 2008; Dio *et al*, 2023; Boden *et al*, 2009; Joyce & McKim, 2010). Based on previous studies and our data, we reason that ASY3 might promote the CO formation in at least two ways. First, as a conserved structural component of the axis and SC, ASY3 facilitates the interhomolog recombination by promoting pairing and synapsis, e.g., via establishing bridges between the recombinational nucleofilament and homolog to promote homology search and strand invasion (Ito & Shinohara, 2023; Dubois *et al*, 2019). This is supported by our data showing that chromosome pairing and synapsis is severely disrupted in *asy3^aacc^*mutants where the axis is likely not formed properly, consistent with previous findings (Figure 4A, 5A) (Ferdous *et al*, 2012). Second, we found that the initial loading of HEI10 on chromosomes is severely disrupted in the absence of ASY3 in both *Arabidopsis* and *Brassica napus* (Figure 6B, Supplemental figure 8B and C). Thus, ASY3 supports the CO formation by ensuring the fidelity of HEI10 loading. Whether other pro-CO factors are also affected by ASY3 remains to be explored.

The question of to which extent the effect of ASY3 dosage reduction on recombination is attributed to its role in ASY1 recruitment remains unclear. Recently, the function of ASY1 was analyzed using a series of tetraploid wheat *asy1* mutants (Dio *et al*, 2023). In contrast to the series of *asy3* allelic mutants reported here, those *asy1* mutants in tetraploid wheat, display a linear reduction in chiasmata (COs) concomitantly with the decrease of *ASY1* gene dosage, resulting in the failure to maintain CO assurance (Dio *et al*, 2023). In those wheat mutants with only one functional allele of ASY1, distal COs prefer to be formed at the expense of proximal and interstitial COs, supporting the conclusion that ASY1 functions to promote CO formation away from the chromosome ends (Dio *et al*, 2023; Lambing *et al*, 2020a). Our data indicate that ASY3 may be an ideal target of manipulation to aim for the global increase of CO frequency in crop breeding without compromising the proximal and interstitial COs.

### Role of ASY3 in HEI10 coarsening and CO interference

Recently, a mechanistic coarsening model that quantitatively explains the class I CO patterning was proposed (Morgan *et al*, 2021; Zhang *et al*, 2021; Fozard *et al*, 2023; Durand *et al*, 2022). According to this model, the SC plays a critical role in controlling and constraining the dynamic coarsening of HEI10 molecules, thus imposing the CO interference. This idea is compatible with the finding that the number of HEI10 foci gets increased in the absence of the transverse filament protein of the SC, ZYP1, where interference is abolished, indicating a role of ZYP1/SC in HEI10 dynamics (Capilla-Pérez *et al*, 2021; France *et al*, 2021). When ASY3 is absent, the initial loading of HEI10 on chromosomes is compromised, resulting in the deficiency of HEI10 and thus reduced CO formation. Notably, HEI10 initial loading appears normal when ASY3 is reduced by ∼ 60-70% in *asy3^aaCc^*. However, this reduction of ASY3 dosage in *asy3^aaCc^* leads to a patchier and less continuous assembly of the SC, which likely causes the HEI10 coarsening to work locally but not along the entire chromosome length, thus compromising the CO interference and increasing CO formation. (France *et al*, 2021)We hence propose that an intact and continuous tripartite structure of the SC is essential for proper HEI10 diffusion and CO interference. In this context, one interesting question is whether the CO formation could be further unleased when combining the manipulation of ASY3/axis and ZYP1.

In conclusion, the results we present here unravel the dosage-dependent diverse effects of ASY3 on CO formation and provide insights into the role of chromosome axis/SC in CO patterning. Since ASY3 is wildly present in a variety of crop species, this work provides a promising target—alone and in combination—for modifying the CO efficiency for crop breeding.

## Materials and methods

### Plant materials and growth condition

The spring-type *Brassica napus* cultivar *Westar* was used as wild-type reference throughout this research. Mutants of *Bnaasy3* were generated by the CRISPR/Cas9 gene editing technique in the background of *Brassica napus* cv. *Westar*. The *Arabidopsis* T-DNA insertion mutants *asy1-4* (SALK_046272) and *hei10-2* (SALK_014624) were previously described (Yang *et al*, 2020b; Chelysheva *et al*, 2012) and used for validating the specificity of ASY1 and HEI10 antibodies used in this study. *Arabidopsis* plants were grown in growth chambers under a cycle of 16 h of light at 21°C and 8 h of dark at 18°C. Rapeseed plants were planted in the experimental fields with normal growing conditions at Huazhong Agricultural University, Wuhan, China.

### Plasmid construction

To generate *Bnaasy3* mutants, two sgRNAs (Figure 1A) were designed to target the first exon of all *ASY3* alleles by CRISPR-P2.0 (http://crispr.hzau.edu.cn/CRISPR2), and then inserted into the binary vector pKSE401 containing the guide RNA and Cas9 expression cassettes via golden gate assembly (Xing *et al*, 2014). For the yeast two-hybrid assay of BnaASY1 with BnaASY3, BnaA07.ASY1^1-300aa^-BD, BnaA05.ASY3^FL^-AD, BnaA05.ASY3^1-32aa^-AD constructs were generated. The CDS sequences of *BnaA07.ASY1* (1-300 aa) and *BnaA05.ASY3* (full length and 1-32aa) were amplified by PCR with primers flanked by attB recombination sites (BnaASY1-A07-attB1F and BnaASY1-A07-300aa-attB2R, BnaASY3-A05-attB1F and BnaASY3-A05-attB2R, BnaASY3-A05-attB1F and BnaASY3-A05-32aa-attB2R) and subcloned into *pDONR223* vector by Gateway BP reactions. All these entry clones were subsequently integrated into the destination vectors *pGADT7-GW* or *pGBKT7-GW* vectors by Gateway LR reactions. Primers used are listed in Supplemental table 3.

### Plant transformation and genotyping

The procedure of agrobacterium-mediated transformation of *Brassica napus* was carried out as previously described using hypocotyl explants (Dai *et al*, 2020). To genotype the CRISPR-Cas9-induced mutations of *ASY3* genes, DNA sequences cover the targeting regions were amplified by PCR using gene-specific primers for *BnaA05.ASY3* and *BnaC04.ASY3* alleles (BnaASY3-A05-F1 and BnaASY3-R1, BnaASY3-C04-F1 and BnaASY3-R1) and were subjected to sequencing. Primers used are listed in Supplemental table 3.

### Yeast two-hybrid assay

For the yeast two-hybrid assay, the relevant combinations of constructs were co-transformed into the auxotrophic yeast strain AH109 using the polyethylene glycol/lithium acetate method according to the manufacturer’s manual (Clontech). Yeast cells were dotted on the plates of double (-Leu-Trp) and quadruple (-Leu-Trp-His-Ade) synthetic dropout medium and images were captured after 3 days of incubation at 28°C.

### Antibody generation

The polyclonal antibodies against Brassica ASY1, ASY3, REC8, and ZYP1 were produced by DIA-AN, Wuhan, China (https://www.dia-an.com). Briefly, the coding regions of BnaC06.ASY1 (301-614 aa), BnaC04.ASY3 (481-696 aa), BnaA10.REC8 (377-701 aa), and BnaA07.ZYP1A (1-349 aa) were amplified and inserted into the pET-32a vector. The corresponding recombinant proteins were produced and purified from *Escherichia coli* bacteria, which were used as antigens to immunize rabbit or rat. After 3-times immunization, antibodies were purified from the antisera using antigen-based affinity purification. The polyclonal antibody against HEI10 was generated by GL Biochem, Shanghai, China (www.glschina.com) using the synthetic peptide (Cys-PANNFYPRHQEP) conjugated to the carrier protein KLH as the antigen.

### Cytological analysis

Pollen viability was performed by dipping the open flowers into the Peterson staining solution (Peterson *et al*, 2010). For tetrad analysis, flower buds at appropriate size were dissected and the resulting anthers were treated in 1mol/L HCl for 1 min at 60°C. Subsequently, anthers were rinsed in distilled water and squashed in cabol fuchsin solution. Images were captured using a SOPTOP ex30 light microscope equipped with a color camera.

Meiotic chromosome spread analysis was performed as described previously with minor modifications (Chelysheva *et al*, 2010). Briefly, fresh flower buds were fixed in the ethanol: acetic acid (3:1) fixative for 24 to 48h at 4°C, then washed twice with the same fixative, and stored at -20°C. For chromosome spreading, flower buds at appropriate size were dissected. Next, anthers were digested in the digestion mix (3% cellulase, 3% macerozyme and 5% snailase in 50 mM citrate buffer, pH 4.5) for 50 min at 37°C. Then, a single digested anther was mashed into fine suspension by a hook in 5μl water on the microscopy slide. Subsequently, 30μl of 60% acetic acid was added to the slide, followed by a gentle stirring without toughing the slide surface using a straight needle for 2 min on a 45°C hotplate. Finally, before the drop dried out, the cells were fixed on the slide by rinsing the slide with cold fixative (ethanol: acetic acid 3:1) and air-drying the preparation. Chromosomes were stained with 4′,6-diamidino-2-phenylindole (DAPI) and observed under the fluorescent microscope equipped with a monochrome camera.

For immunostaining, following the preparation of chromosome spreads, the slides were put into a glass jar filled with 10 mM citrate buffer pH 6.0, microwaved until slight boiling, and then transferred immediately into 1x PBST solution (0.1% Triton X-100). Next, the slides were first blocked with goat serum for 1-2 h at 28°C and then incubated with relevant antibodies for 48h at 4°C. After three times of washing (5 mins each) in PBST, the slides were incubated with fluorescein-conjugated secondary antibodies (ThermoFisher) for 24 h at 4°C. Finally, the slides were washed twice (5 mins each) with PBST and stained by DAPI. Images were captured using the fluorescent microscope equipped with a monochrome camera.

### Quantification of the fluorescent intensity and relative distance of HEI10 foci

To quantify the signal intensity of ASY3 and ASY1, three to five slides were prepared in the same batch of experiment for each genotype using flower buds from different plants, which were then treated in parallel under same conditions to minimize the slide-to-slide variation, e.g., the same amount and duration of antibody incubation followed by same strength of washing steps. All images were captured under the same exposure conditions. 10 to 15 cells were recorded from each slide. The signal intensity was measured by Fiji. The background signal was subtracted. All raw values were normalized through dividing by the mean signal intensity value of wildtype. Noting that for the quantification of ASY1 signal intensity, the “blob”-like overexposed regions (red arrowheads in Figure 2, Supplemental figure 4A) were cut out.

For the quantification of the relative distance of adjacent HEI10 foci on each bivalent, the distance between two adjacent HEI10 foci on a single bivalent were measured using Fiji and then normalized by the total length of the bivalent.

### Statistical analysis

The one-way ANOVA followed by Game-Howell’s or Tukey’s multiple comparisons test and students’ t-test were performed using GraphPad Prism 8.0.2 software. For multiple comparisons test, the Tukey’s multiple comparison test is applied when the variances of different groups of data are equal (by F-test), and otherwise, the Game-Howell’s multiple comparisons test is used. The calculation of the mean and standard deviation, the Poisson distribution comparison, and the Chi-square test were done using Microsoft Excel.

## Acknowledgements

We thank Dr. Maren Heese and Dr. Joke De Jaeger Braet (both from University of Hamburg) for the critical reading of the manuscript. This work was supported by the Scientific Innovation 2030 Project from China (2022ZD0401004), National Natural Science Foundation of China (32170354, 32370360), and Fundamental Research Funds for the Central Universities (2662023ZKPY003 and 2662023PY004).

## Author contributions

C.Y. and B.Y. conceived this research. L.C. performed most of the experiments. J.Z., M.G., and Y.Z. contributed to the genetic transformation, plant growing, and chromosome spreading. L.C., C.Y., B.Y., and C.Z. analyzed the data. C.Y., A.S., and L.C. wrote the manuscript.

---

**Supplemental figure 1.**
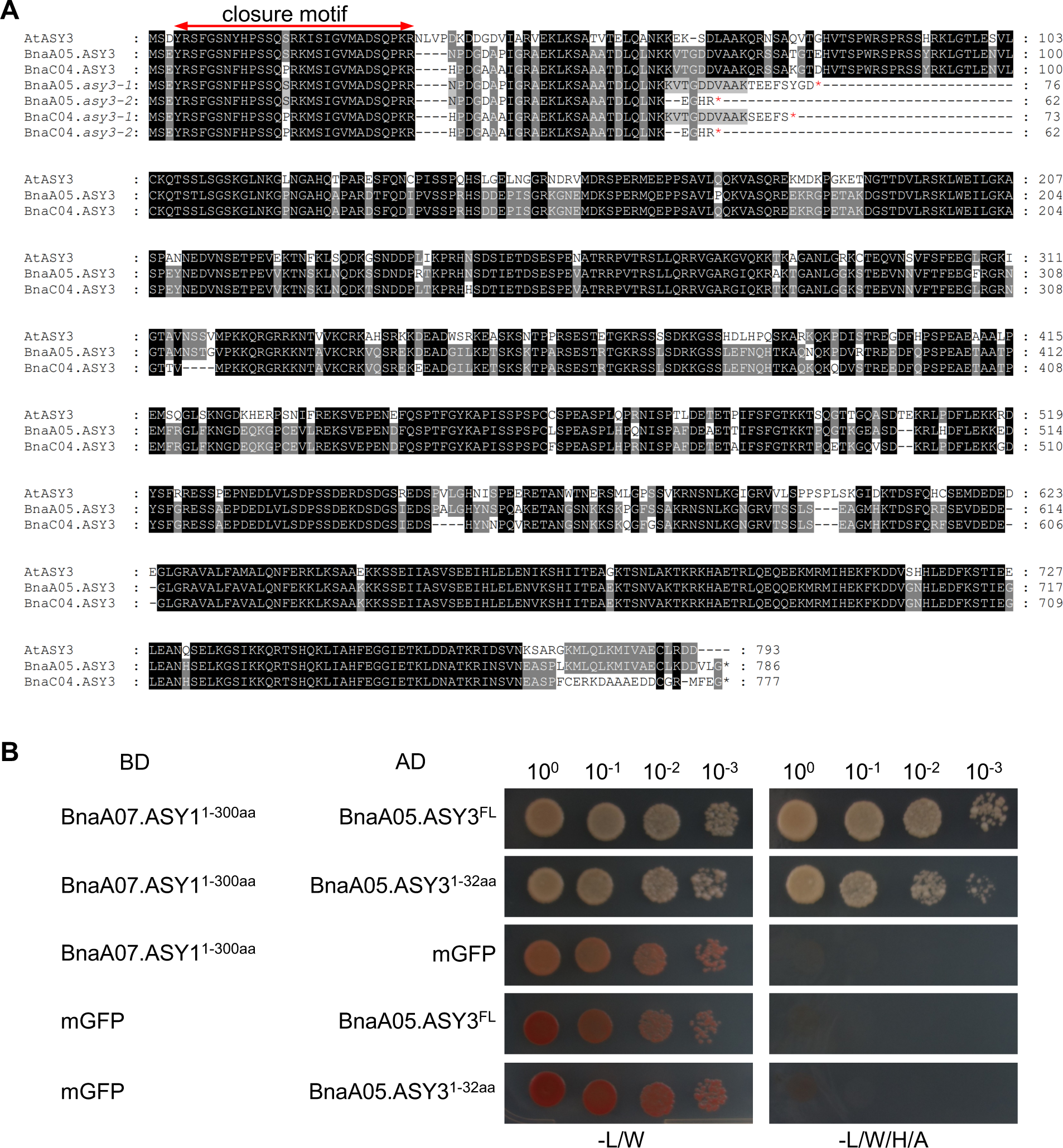
Analysis of ASY3 in *Brassica napus*. (A) Alignment of ASY3 proteins from *Arabidopsis* and *Brassica napus*. Red asterisks indicate the putative translational stops of mutated ASY3 proteins. The region of the closure motif is highlighted by the red line with double arrowheads. (B) Yeast two-hybrid assay testing for interaction of BnaA07.ASY1 HORMA domain (1-300 aa) with BnaA05.ASY3 closure motif (1-32aa). Yeast cells harbouring both the AD and BD plasmids were grown on synthetic dropout (SD) medium in the absence of Leu and Trp (-L/W, left panel) and of Leu, Trp, His, and Ade (-L/W/H/A, right panel). Yeast cells were incubated until OD_600_=1 and then diluted 10-, 100-, and 1,000-fold for the assay.

**Supplemental figure 2.**
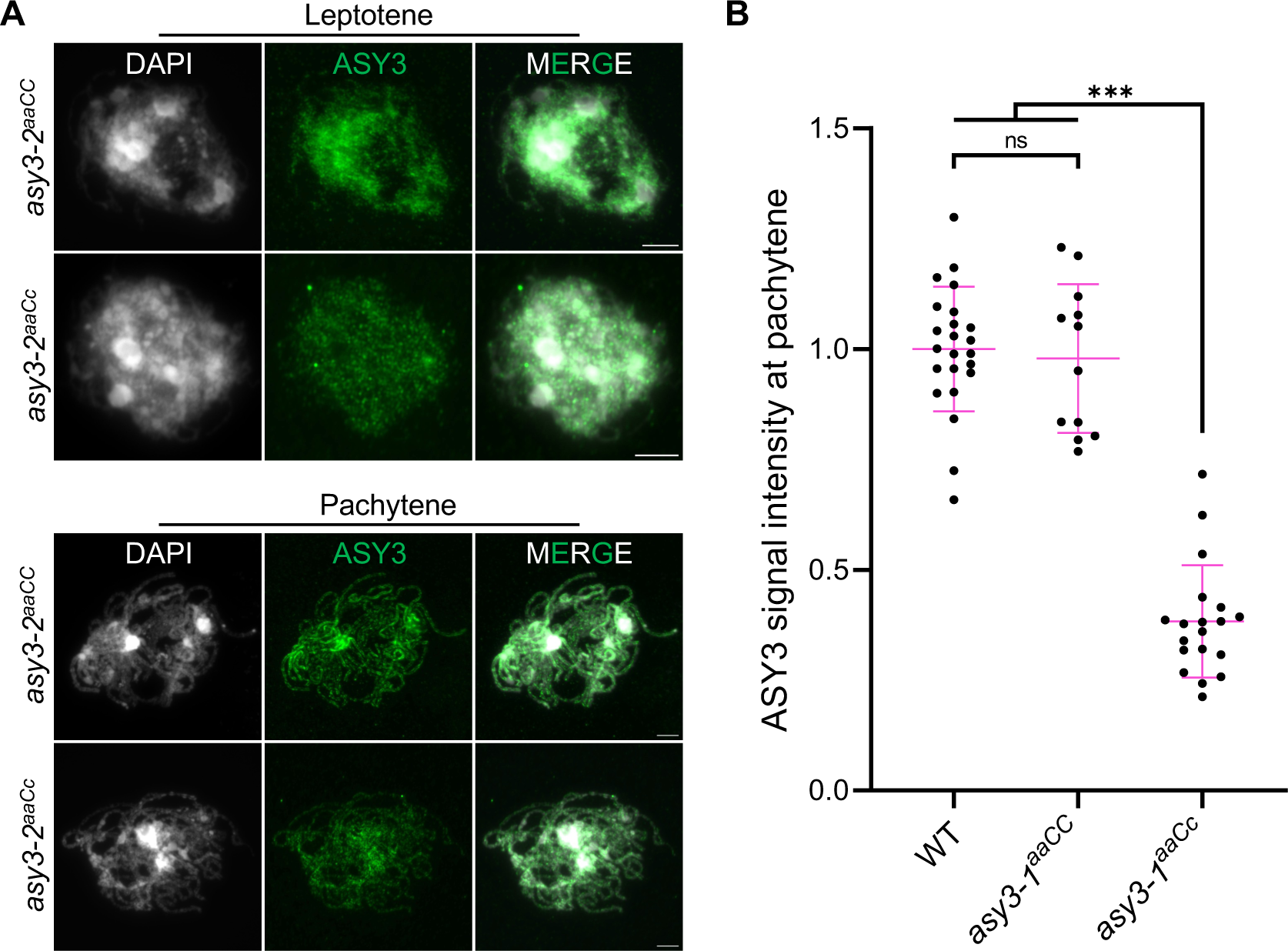
Analysis of ASY3 localization in *asy3^aaCC^* and *asy3^aaCc^* mutants. (A) Immunolocalization of ASY3 in male meiocytes of *asy3-2^aaCC^* and *asy3-2^aaCc^* mutants at leptotene and pachytene (or -like). Bars: 5µm. (B) Quantification of relative ASY3 signal intensity at pachytene in WT, *asy3-1^aaCC^*, and *asy3-1^aaCc^* mutants. Asterisks indicate significant difference (Game-Howell’s multiple comparisons test, *p*<0.001).

**Supplemental figure 3.**
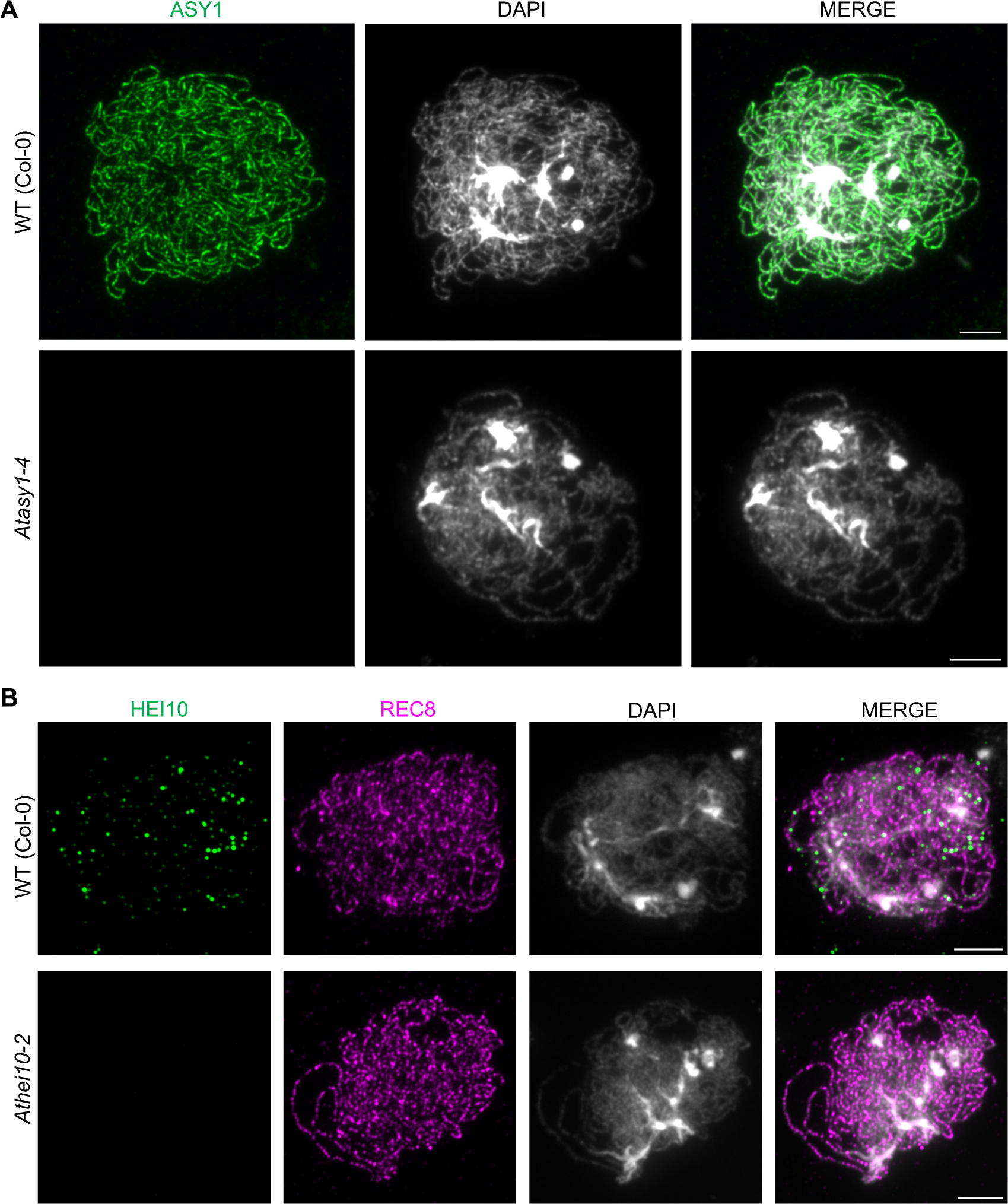
Specificity validation of ASY1 and HEI10 antibodies used in this study. (A) Immunostaining of ASY1 in male meiocytes of *Arabidopsis asy1-4* mutant at early prophase I. (B) Co-immunostaining of HEI10 and REC8 in male meiocytes of *Arabidopsis hei10-2* mutant at early prophase I. Bars: 5 µm.

**Supplemental figure 4.**
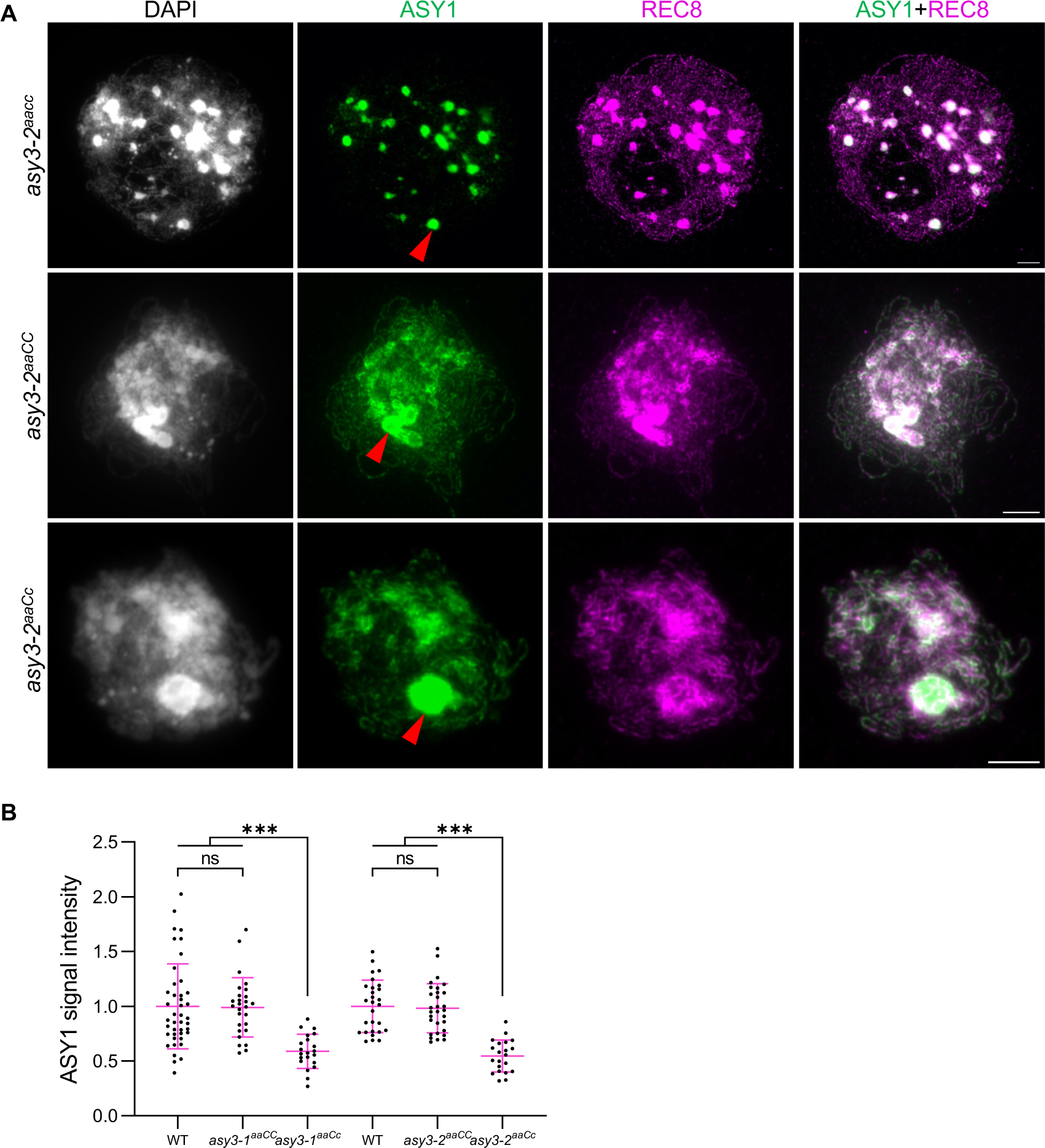
Localization of ASY1 in *asy3* null and hypomorphic mutants. (A) Co-immunostaining of ASY1 and REC8 in male meiocytes of *asy3-2^aacc^*, *asy3-2^aaCC^*, and *asy3-2^aaCc^* mutants at leptotene. Red arrowheads indicates the “blob”-like regions with overexposed signal. (B) Relative ASY1 signal intensity in WT, *asy3-1^aaCC^*, *asy3-1^aaCc^*, *asy3-2^aaCC^*, and *asy3-2^aaCc^* mutant plants at leptotene. The comparisons of signal intensity of WT, *asy3-1^aaCC^*, and *asy3-1^aaCc^* and WT, *asy3-2^aaCC^*, and *asy3-2^aaCc^* mutant plants were plotted independently. Noting that the “blob”-like overexposed regions (red arrowheads) were removed from the quantification. Asterisks indicate significant difference (Game-Howell’s multiple comparisons test, *p*<0.001).

**Supplemental figure 5.**
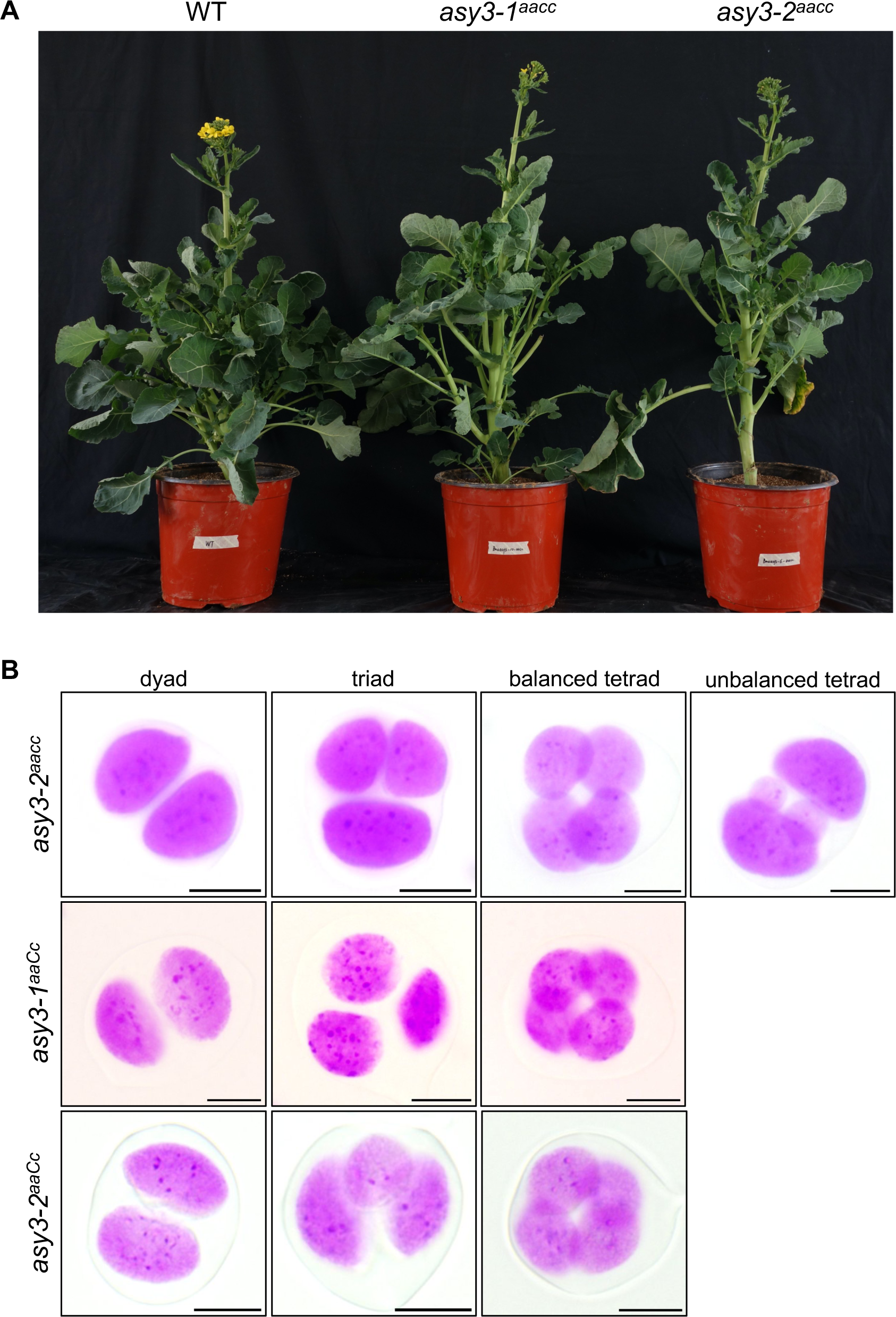
Phenotypic analysis of *asy3* mutants. (A) Plants of WT, *asy3-1^aacc^*, and *asy3-2^aacc^* mutants at early flowering stage. (B) Representative images of male meiotic products in *asy3-2^aacc^*, *asy3-1^aaCc^*, and *asy3-2^aaCc^* mutant plants. Bars: 5µm.

**Supplemental figure 6.**
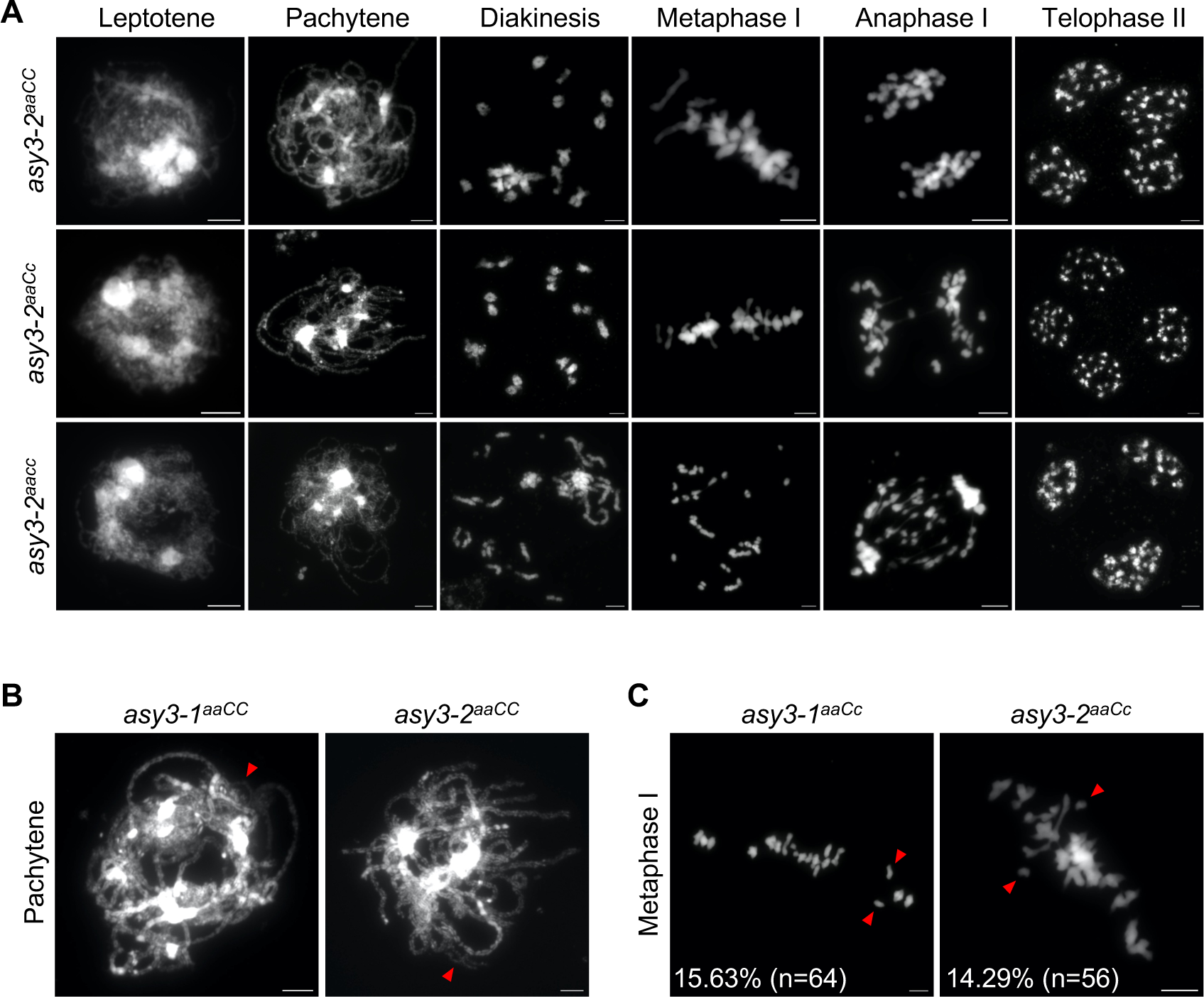
Analysis of meiotic chromosome behaviors in *asy3* mutants. (A) Chromosome spread analysis of male meiosis in *asy3-2^aacc^*, *asy3-2^aaCC^*, and *asy3-2^aaCc^* mutants. (B) Representative images showing the unpaired stretches (red arrowheads) in male meiocytes of *asy3-1^aaCC^*and *asy3-2^aaCC^* at pachytene. (C) Representative cells having one pair of univalents (red arrowheads) in male meiocytes of *asy3-1^aaCc^* and *asy3-2^aaCc^* at metaphase I. The percentages indicate the ratio of cells with one pair of univalents. Bars: 5µm.

**Supplemental figure 7.**
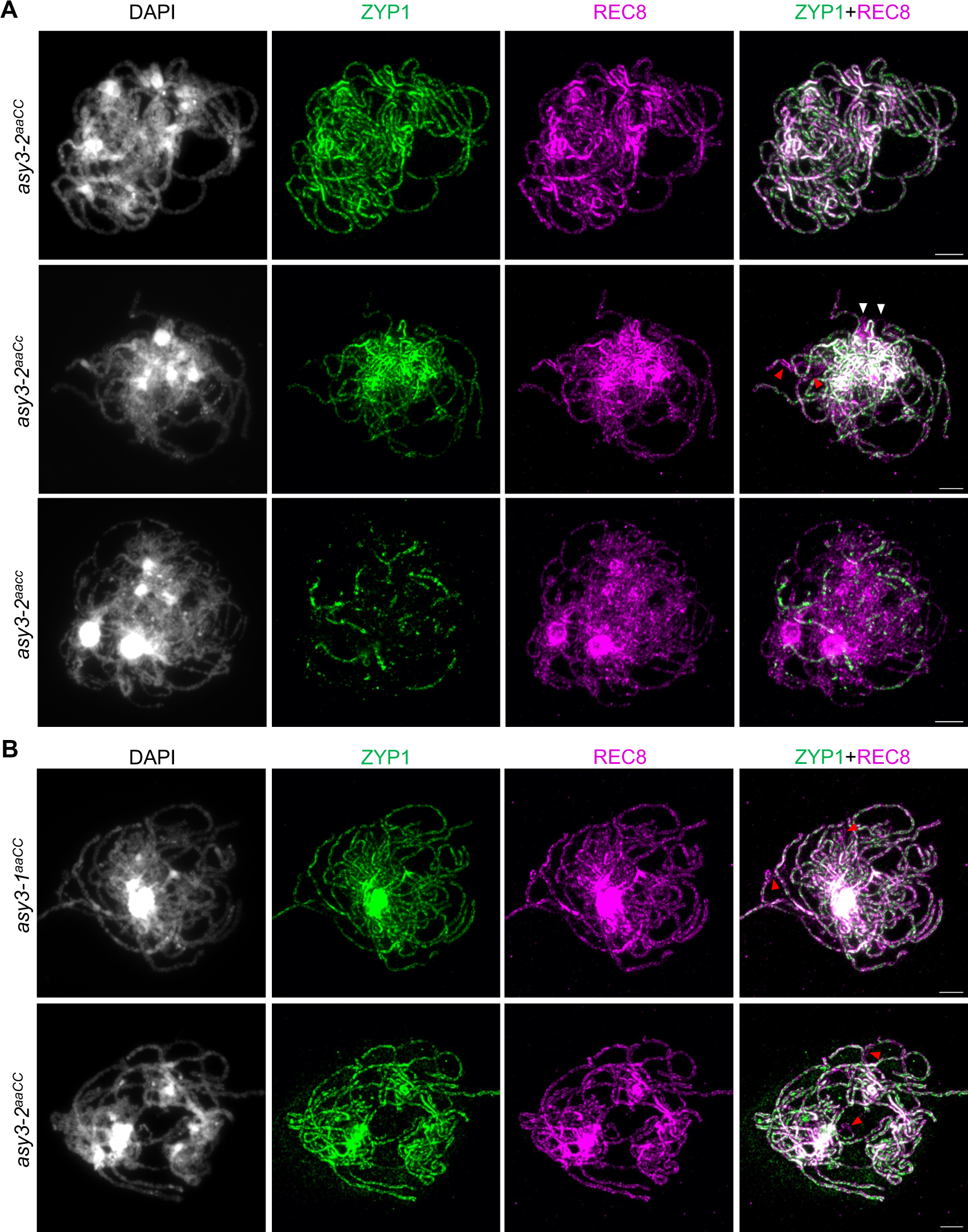
ASY3 dosage-dependent effects on chromosome synapsis. (A) Co-immunostaining of ZYP1 and REC8 in male meiocytes of *asy3-2^aaCC^*, *asy3-2^aaCc^*, and *asy3-2^aacc^* mutants at pachytene. White and red arrowheads indicate the unpaired single threads or coaligned regions that both have no ZYP signal, respectively. (B) Representative images show cells having a bit ZYP1 non-labeled regions in male meiocytes of *asy3-1^aaCC^* and *asy3-2^aaCC^* at pachytene. Red arrowheads indicate non-ZYP1 labeled regions. Bars: 5µm.

**Supplemental figure 8.**
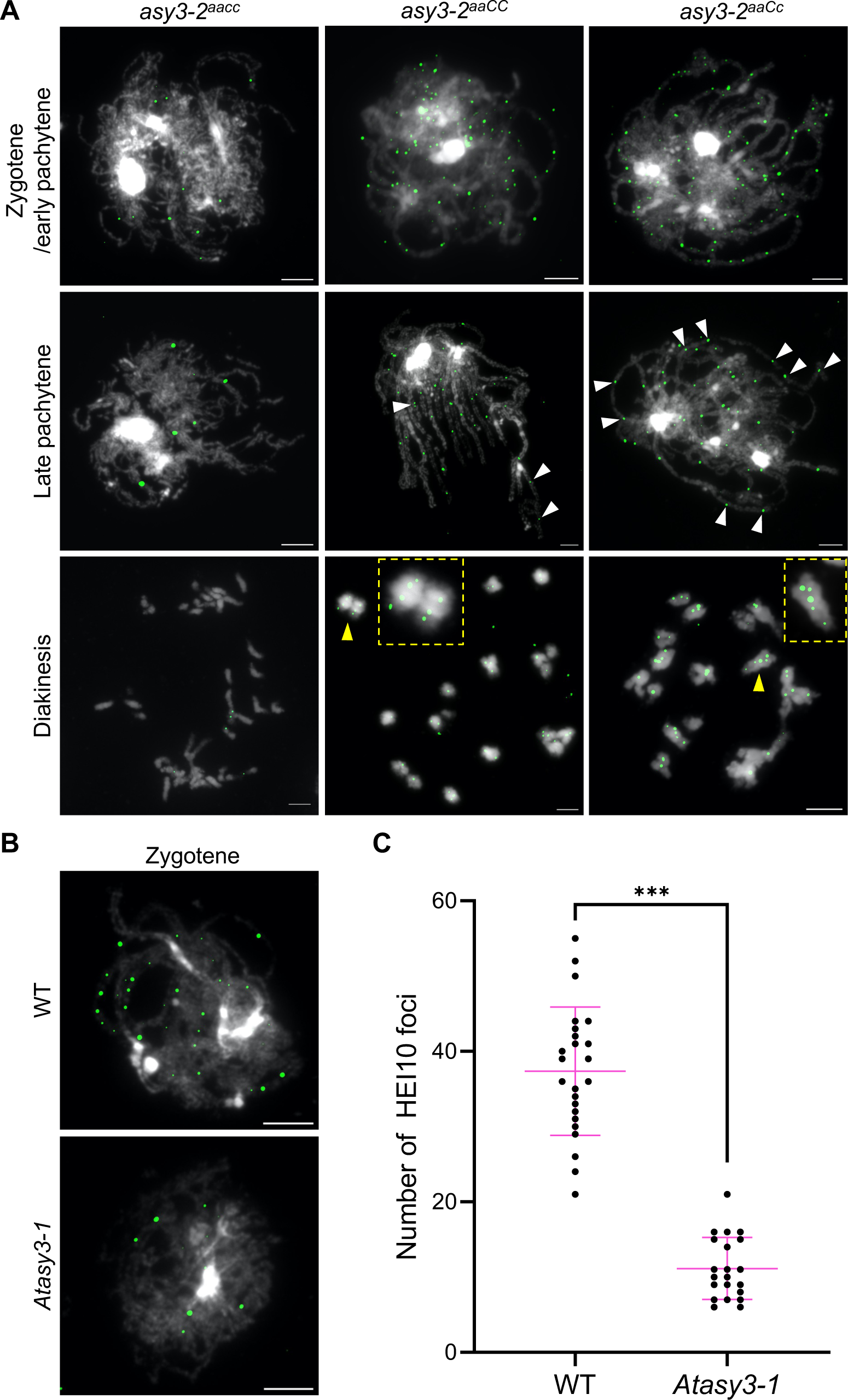
Analysis of HEI10 localization in *asy3* mutants of *Arabidopsis* and *Brassica napus*. (A) Immunolocalization of HEI10 in male meiocytes of *asy3-2^aacc^*, *asy3-2^aaCC^*, and *asy3-2^aaCc^* mutants at zygotene/early pachytene, late pachytene, and diakinesis. White arrowheads indicate closely localized HEI10 foci alone one chromosome pair. Yellow arrowheads depict the magnified bivalents shown in the yellow rectangles. Bars: 5µm. (B) Immunolocalization of HEI10 at early prophase I in male meiocytes of Arabidopsis WT and *asy3-1* mutants. Bars: 5µm. (C) Quantification of the number of HEI10 foci in Arabidopsis WT and *asy3-1* mutants at early prophase I (zygotene/early pachytene). The statistical analysis was performed by student’s t-test (*** *p*<0.001).

**Supplemental table 1.** Summary of chromosome configurations at diakinesis/metaphase I in male meiocytes of wildtype and *asy3* mutants. Values represent the mean±SD followed by the percentage.

| Genotype | Ring bivalent | Rod bivalent | Univalent | Number of PMCs |
| --- | --- | --- | --- | --- |
| Wild type | 12.53±1.56 (65.95%)<sup>b</sup> | 6.47±1.57 (34.05%)<sup>a</sup> | 0.00±0.00 (0.00%) | 32 |
| *asy3-1<sup>aacc</sup>* | 0.27±0.53 (1.40%)<sup>c</sup> | 4.88±1.33 (25.67%)<sup>b</sup> | 27.71±2.80 (72.93%)<sup>a</sup> | 49 |
| *asy3-2<sup>aacc</sup>* | 0.21±0.41 (1.11%)<sup>c</sup> | 5.74±1.97 (30.19%)<sup>b</sup> | 26.11±3.46 (68.70%)<sup>a</sup> | 19 |
| *asy3-1<sup>aaCc</sup>* | 15.57±1.41 (81.92%)<sup>a</sup> | 3.35±1.43 (17.62%)<sup>c</sup> | 0.17±1.56 (0.46%)<sup>b</sup> | 23 |
| *asy3-2<sup>aaCc</sup>* | 15.00±1.83 (78.95%)<sup>a</sup> | 3.93±1.84 (20.70%)<sup>bc</sup> | 0.13±0.50 (0.35%)<sup>b</sup> | 15 |
| *asy3-1<sup>aaCC</sup>* | 14.06±1.72 (73.98%)<sup>ab</sup> | 4.89±1.70 (25.73%)<sup>bc</sup> | 0.11±0.46 (0.29%)<sup>b</sup> | 18 |
| *asy3-2<sup>aaCC</sup>* | 12.88±2.09 (67.76%)<sup>ab</sup> | 6.00±2.24 (31.58%)<sup>ab</sup> | 0.25±0.66 (0.66%)<sup>b</sup> | 8 |

**Supplemental table 2.**
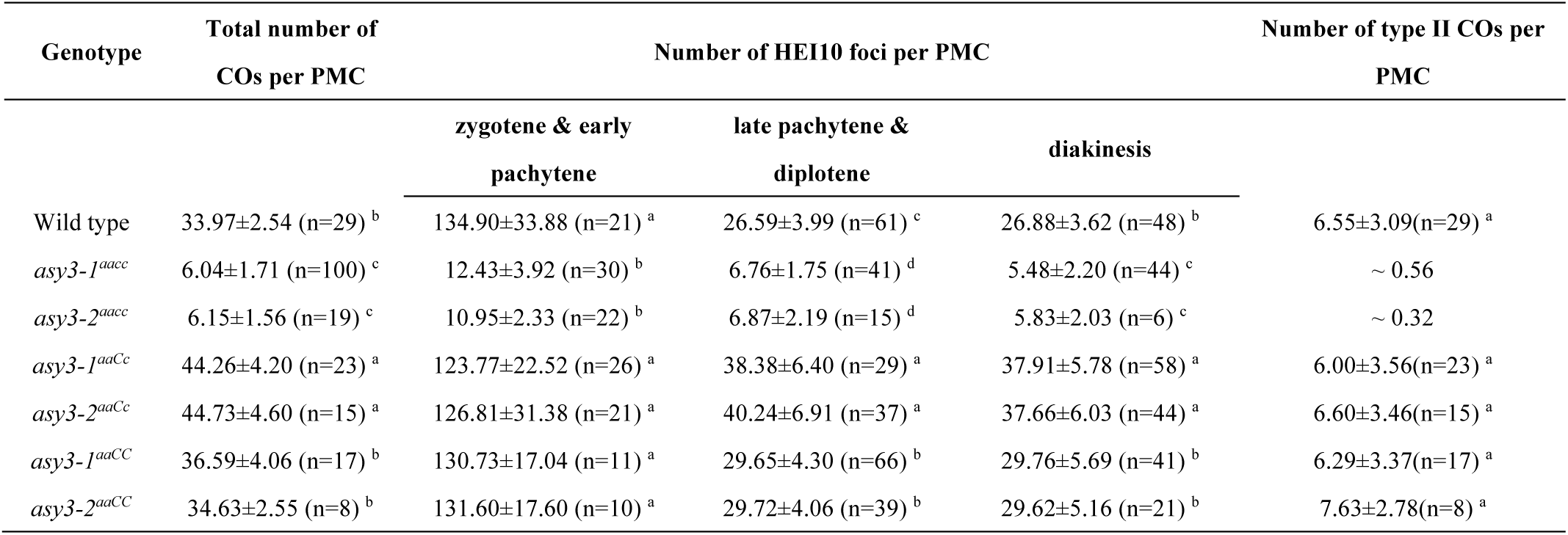
Summary of the number of HEI10 foci, total number of COs, and number of type II COs in male meiocytes of wildtype and *asy3* mutants. Values represent the mean±SD followed by the number of observed nuclei.

| Genotype | Total number of COs per PMC | Number of HEI10 foci per PMC |  |  | Number of type II COs per PMC |
| --- | --- | --- | --- | --- | --- |
|  |  | zygotene & early pachytene | late pachytene & diplotene | diakinesis |  |
| Wild type | 33.97±2.54 (n=29)<sup>b</sup> | 134.90±33.88 (n=21)<sup>a</sup> | 26.59±3.99 (n=61)<sup>c</sup> | 26.88±3.62 (n=48)<sup>b</sup> | 6.55±3.09(n=29)<sup>a</sup> |
| *asy3-1<sup>aacc</sup>* | 6.04±1.71 (n=100)<sup>c</sup> | 12.43±3.92 (n=30)<sup>b</sup> | 6.76±1.75 (n=41)<sup>d</sup> | 5.48±2.20 (n=44)<sup>c</sup> | ~ 0.56 |
| *asy3-2<sup>aacc</sup>* | 6.15±1.56 (n=19)<sup>c</sup> | 10.95±2.33 (n=22)<sup>b</sup> | 6.87±2.19 (n=15)<sup>d</sup> | 5.83±2.03 (n=6)<sup>c</sup> | ~ 0.32 |
| *asy3-1<sup>aaCc</sup>* | 44.26±4.20 (n=23)<sup>a</sup> | 123.77±22.52 (n=26)<sup>a</sup> | 38.38±6.40 (n=29)<sup>a</sup> | 37.91±5.78 (n=58)<sup>a</sup> | 6.00±3.56(n=23)<sup>a</sup> |
| *asy3-2<sup>aaCc</sup>* | 44.73±4.60 (n=15)<sup>a</sup> | 126.81±31.38 (n=21)<sup>a</sup> | 40.24±6.91 (n=37)<sup>a</sup> | 37.66±6.03 (n=44)<sup>a</sup> | 6.60±3.46(n=15)<sup>a</sup> |
| *asy3-1<sup>aaCC</sup>* | 36.59±4.06 (n=17)<sup>b</sup> | 130.73±17.04 (n=11)<sup>a</sup> | 29.65±4.30 (n=66)<sup>b</sup> | 29.76±5.69 (n=41)<sup>b</sup> | 6.29±3.37(n=17)<sup>a</sup> |
| *asy3-2<sup>aaCC</sup>* | 34.63±2.55 (n=8)<sup>b</sup> | 131.60±17.60 (n=10)<sup>a</sup> | 29.72±4.06 (n=39)<sup>b</sup> | 29.62±5.16 (n=21)<sup>b</sup> | 7.63±2.78(n=8)<sup>a</sup> |

**Supplemental table 3.**
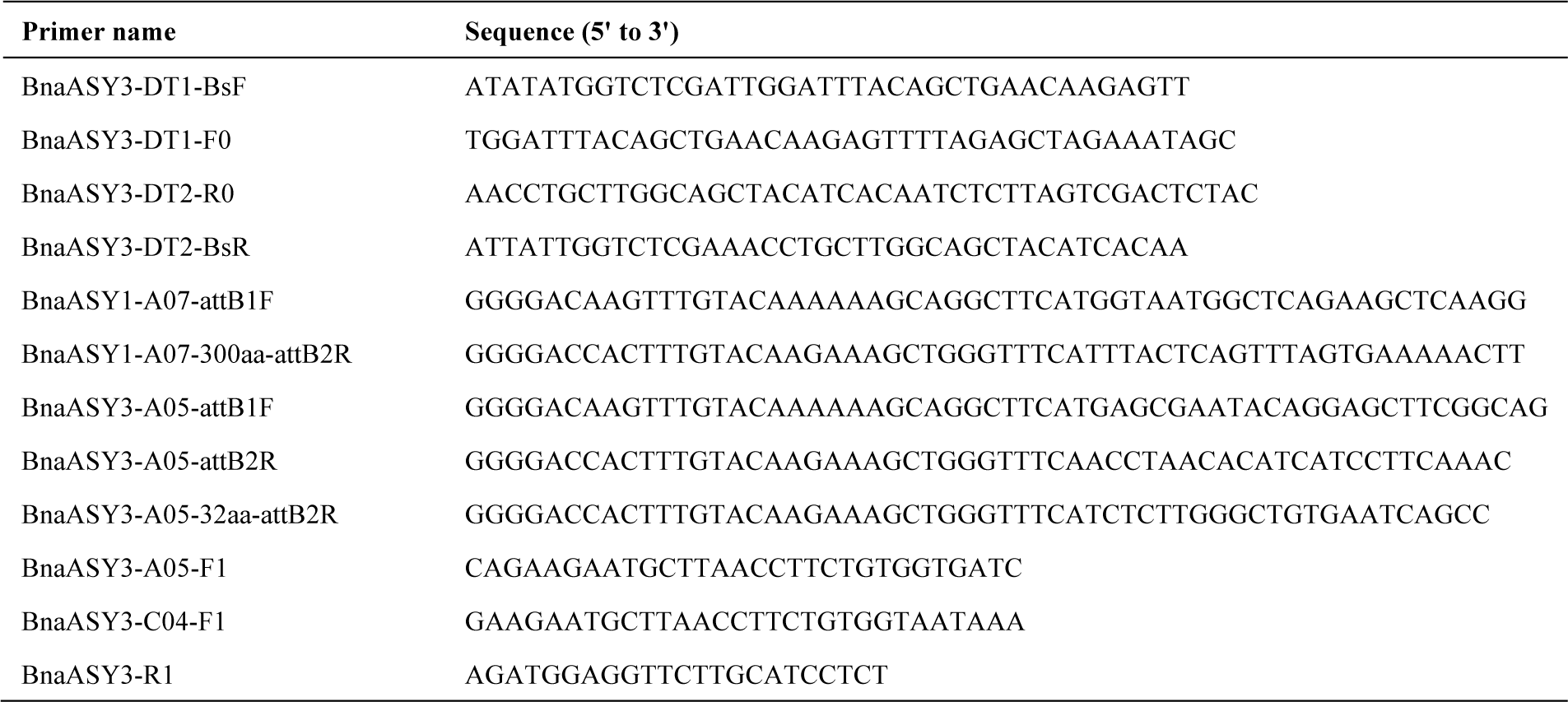
Primers used in this research.

